# Constructing local Cell Sepcific Networks from Single Cell Data

**DOI:** 10.1101/2021.02.13.431104

**Authors:** Xuran Wang, David Choi, Kathryn Roeder

## Abstract

Single-cell RNA sequencing facilitates investigations of gene co-expression networks at the cellular level potentially yielding unique biological insights, but the estimation problem is challenging. We develop a non-parametric approach to estimate cell-specific networks for each cell and cell type and propose novel downstream analyses. The individual networks preserve the cellular heterogeneity and facilitate testing for differences between cell groups. To further our understanding of autism spectrum disorder, we examine the evolution of gene networks in fetal brain cells and compare the gene networks of cells sampled from case and control subjects to reveal patterns in gene co-expression.

## Background

Single-cell RNA-sequencing (scRNA-seq) provides a high throughput profile of RNA expression for individual cells that reveals the heterogeneity across cell populations. Recent advances in computational methods enable cell-type classification, novel cell-type identification (39), and trajectory alignment (30); however, among these methods, less attention has been devoted to gene-gene association and transcriptional networks, which can shed light on many vital biological processes. Gene expression is known to be tightly regulated by networks of transcription factors and understanding these networks at a cellular level can identify differences in the inner workings of cells in normal and diseased tissues (10, 40, 28, 11).

Thus far, efforts to estimate co-expression networks from scRNA-seq have been limited in their success (12, 35) for several reasons, including technical challenges such as a sparsity of non-zero counts and a high level of noise (5), nonlinear relationships that are not easily captured by traditional measures, and heterogeneity in co-expression patterns across cells.

Efforts have been made to circumvent these challenges (40, 28, 21, 22, 14). Many approaches aim to estimate a single co-expression network across all cell-types, but to gain access to the subtle differences in gene co-expression at the cellular level we require estimates of gene networks evaluated for each cell-type. Moreover, even if the aim is to estimate a cell-type specific network, traditional methods estimate a single co-expression relationship across the entire sample of cells of that type. With such an approach heterogeneity of co-expression across individual cells is erased, but this knowledge can provide valuable insights for downstream analysis.

To reveal co-expression networks within individual cells, we aim to construct cell-specific networks (CSNs) on the cellular level from scRNA-seq data. This idea was first proposed by Dai et al.(7) and Li et al.(20), wherein they generate individual transcriptional networks at a single-cell level for the first time. The construction of CSN is based on the following assessment of local statistical dependency of a pair of genes: for an individual cell of a particular type, if the co-expression of the pair of genes is unusually high, relative to their distribution assuming the genes to be independent, then we infer a gene-gene relationship (Figure 1). This type of analysis produces a full gene-gene network for each cell that can be envisioned as a graph, with genes as nodes and edges depicting gene-gene dependencies. For each cell, Dai et al.(7) and Li et al.(20) then record the degree sequence of its cell-specific network, to be used as input for clustering or low-dimensional embedding.

**Figure 1:**
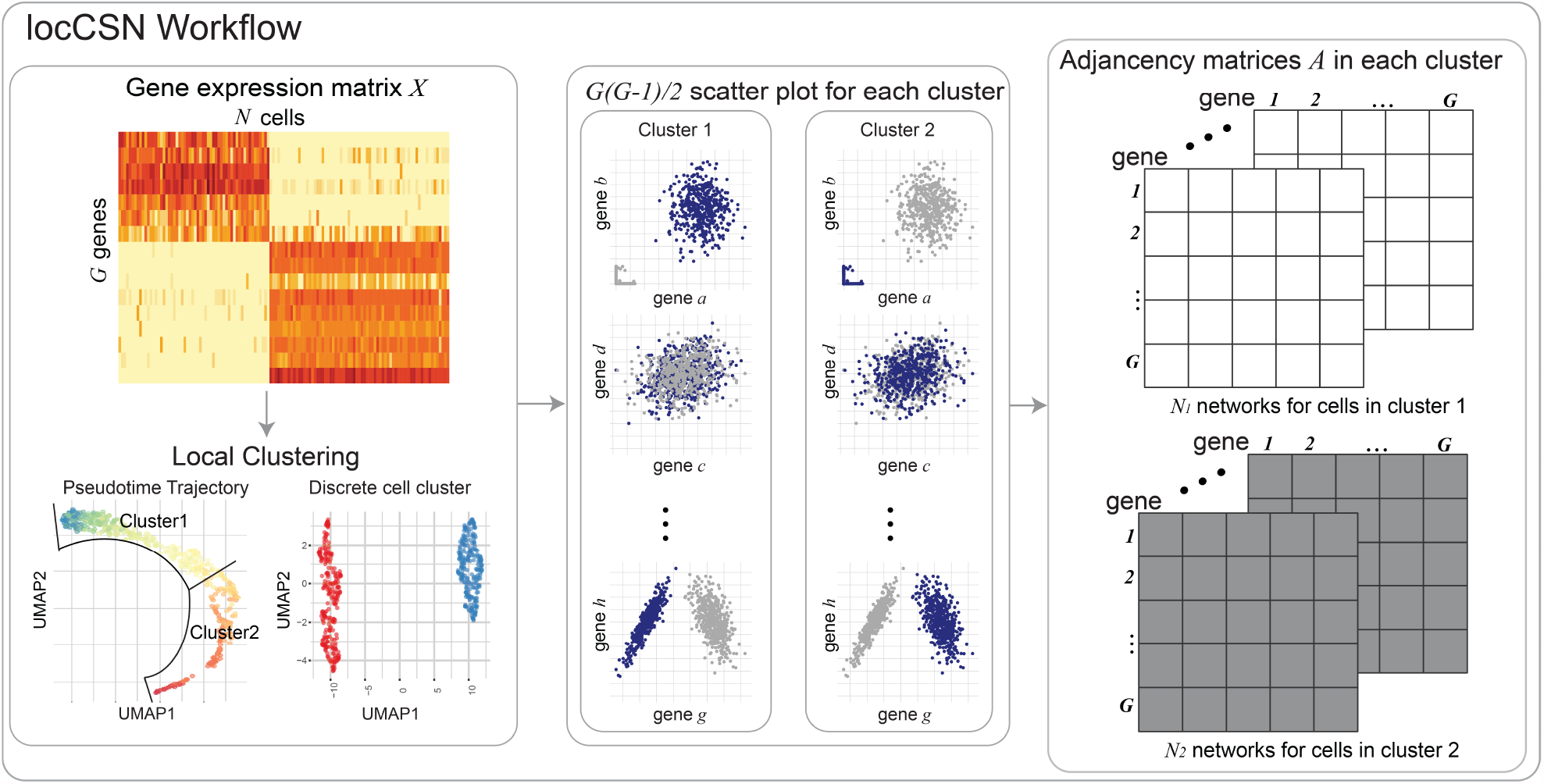
Workflow of locCSN. Starting with the gene expression matrix, we cluster homogeneous cells. For each cell in a cluster, we examine the co-expression of each pair of genes to determine if the joint expression is unusually dense in a neighborhood of the cell, relative to expectation assuming the genes are independent; if so, the genes are connected in that cell’s network. For each cell, we then construct the gene-gene network, based on the results of the local independence test for each pair of genes.

We propose an alternative analysis scheme that expands the utility of the CSN concept for gaining insights into cell biology. We retain the individual networks, rather than just their degree sequences, so that differences or developmental changes can be examined at the level of the gene-gene co-expressions. The individual networks can be used in the following ways: First, averaging CSNs across cells within a category or developmental period provides a nonparametric estimator of the gene network that performs well for the sparse counts generally observed for scRNA-seq data, for example allowing for estimation of evolving gene communities in developmental trajectories. Second, having a sample of CSNs provides a valuable measure of the variability across cells that can be leveraged in powerful tests of network differences across cell groups. For example, comparisons can be made between cells obtained from case and control subjects, or cells sampled from different spatial or temporal regimes. Finally, the CSN approach facilitates followup analysis to discover the key genes driving the differences between networks. With this approach we can identify ‘differential network (DN) genes’, which do not differ in gene expression, but do differ in terms of the co-expression network. These genes could play a key role in network regulation and hence differences that might help explain etiology of disease.

These applications will require more accurate estimates of the individual network connections; we accomplish this by modifying the original network construction algorithm to produce a more adaptive and powerful method, which we call locCSN. We illustrate the value of locCSN by applying it to several scRNA-seq datasets, including developing fetal brains (27), and case and control samples from autism spectrum disorder (ASD) subjects (37). locCSN extracts cell level network information from these data which preserve cell-level network heterogeneity and highlights network differences between case and control samples shedding light on important functional differences.

## Results

Our results are summarised as follows. First we describe the locCSN algorithm, Second, simulated and real data examples are used to illustrate differences between locCSN and oCSN. Then, we apply time-varying community detection to CSNs taken from two overlapping developmental trajectories in the developing cortex atlas dataset, and identify differences in their community evolution. Finally, we show that CSNs enable better separation of case and control populations in the ASD brain dataset, by identifying genes that have differing co-expression (but similar expression levels) between the two groups.

### CSN construction

The algorithm for construction of CSN as originally published (7) (oCSN) is to estimate for each cell a connection strength between each pair of genes, resulting a gene-gene network for every cell. This connection strength is based on a local independence test that is applied to each gene pair and each cell. To motivate this construction, Dai et al.(7) give examples where the test succeeds in separating a mixture of cells, where genes *x* and *y* follow a dependent relationship in some cells but are independent in others (Additional file 1: Figure S2**a**). However, the test requires a choice of resolution (i.e., a bandwidth or window size), which greatly affects performance. Dai et al.(7) proposed a fixed quantile range for the window size, but we find that this approach gives counter-intuitive results and low power when the joint distribution of genes *x* and *y* is a simple correlated normal (Additional file 1: Supplementary Note 1 and Figure S1 - S2). To improve performance, we propose a new local algorithm for CSN construction (locCSN) that allows for the window size of the local independence test to vary cell by cell.

### Usage

While Dai et al.(7) uses all expressions of all cell-types to construct each cell-specific network, we recommend that each cell-specific network be constructed using only the cells of a common type, so that the independence test is conditional on cell type. Otherwise, false edges may be detected due to Simpson’s paradox: given a pair of marker genes that display independent expression within the marked cell-type, CSN construction on all cell-types together will likely infer an edge indicating non-independence of the genes due to the differential expression across types. Therefore we recommend clustering the cells into distinct cell types, and then applying locCSN separately to each cluster, as illustrated for a single cluster in Figure 1. Similarly, if cells are developing along a smooth trajectory, then the cells can be windowed according to pseudotime (30), and each CSN can be computed using the cells within its psuedotime bin.

For each cell, locCSN produces a test statistics for each pair of genes, *z*_*xy*_, to evaluate pairwise gene-gene independence in the neighborhood of that cell’s expression. By thresholding these tests we obtain a 0-1 adjacency matrix for each individual cell. Since *z*_*xy*_ is approximately normally distributed, a natural threshold is the standard normal quantile, *Z*_(1*-α*)_ for *α* chosen to correspond with a natural choice for significance testing, such as *α* = 0.05 or 0.01. The corresponding entry in the adjacency matrix is *I* (*z*_*xy*_ *> Z*_(1*-α*)_), which is either 0 or 1. An estimate of the network for a cluster of cells such as a cell-type is obtained by averaging the adjacency matrices. This average detects unusual clustering of co-expression between genes and thus provides a natural nonparametric estimator of the dependencies in gene expression. The measure is positive by design.

Implementation of locCSN requires two key choices: an initialization of the window width for each local test and the thresholding parameter *α* to derive the 0-1 adjacency matrix from the matrix of local test statistics. We simulate coexpression networks using ESCO (36) to evaluate performance of network estimation for various choices of these tuning parameters. We observe that the standard deviation approach for window width recommended by locCSN performs markedly better than the quantile approach utilized by oCSN. The performance is robust to choice of starting value for the standard deviation algorithm (Additional file 1: Figure S3 - S6). Our results also show that there is a range of threshold *α* for which performance is stable and the optimal choice varies somewhat depending on the strength of the true correlations. Nevertheless, for both moderate and strong correlations performance is good between 0.05 and 0.01 after which the accuracy drops precipitously (Additional file 1: Figure S5 - S6).

### Illustrative Examples

To illustrate the importance of local calculations we compare locCSN and oCSN for synthetic data from ESCO (36). The cells are sampled from two populations, one for which a set of genes exhibit pairwise correlation and another for which the genes are independent (Additional file 1: Figure S8**a** - **c**). When calculated for this set of genes, ideally the test statistics for each cell will distinguish the pattern of correlation present in one population (Group2) not in the other (Group1) (Additional file 1: Figure S8**c**). The desired pattern is achieved by locCSN, but oCSN produces false signals of correlation for many cells in Group1 (Additional file 1: Figure S8**d** and **e**). Mixing the two populations leads to false positives in Group1 suggesting that the false positives result from Simpson’s paradox. Likewise simulations show that BigSCale (14), a powerful alternative approach for coexpression estimation, yields many false connections not detected by locCSN (Additional file 1: Figure S8**f**).

To demonstrate and evaluate the performance of CSN test statistics on real data, we use the Chutype dataset (6). The Chutype dataset includes 7 cell-types (Figure 2**a**). Marker genes corresponding to developmental lineage are provided by the authors and a heatmap of gene expression reveals that a subset of genes mark cell-types DEC and NPC fairly well (Figure 2**b**). We analyze these two cell-types, which contain 138 cells and 173 cells, respectively. The absolute Pearson’s correlation for lineage marker genes, computed within DEC and NPC cell-types, do not show a clear pattern; specifically, the correlation does not delineate the expected block structure for marker genes for DEC and NPC cells (Figure 2**c**). By contrast, averaged locCSNs, thresholded at *α* = 0.05, preserve the gene block structures and emphasizes the differences between cell-types (Figure 2**d**). We conclude that CSN works well on distinct cell-types. The averaged CSN preserves gene blocks, distinguishes between cell-types, and depicts a clearer co-expression pattern than Pearson’s correlation. By contrast, the averaged CSNs calculated from oCSN ignore co-expression between genes, especially the dense block for NPC at the upper right corner (Additional file 1: Figure S9). The performance advantage of locCSN relative to Pearson’s is supported by simulations (Additional file 1: Figure S7 and Table S1). These results are notable because the simulation generates a linear relationship between correlated genes. Thus even when the correlation between genes follows the assumed parametric form, locCSN performs better because it adapts to the sparsity expected in scRNA-seq data.

**Figure 2:**
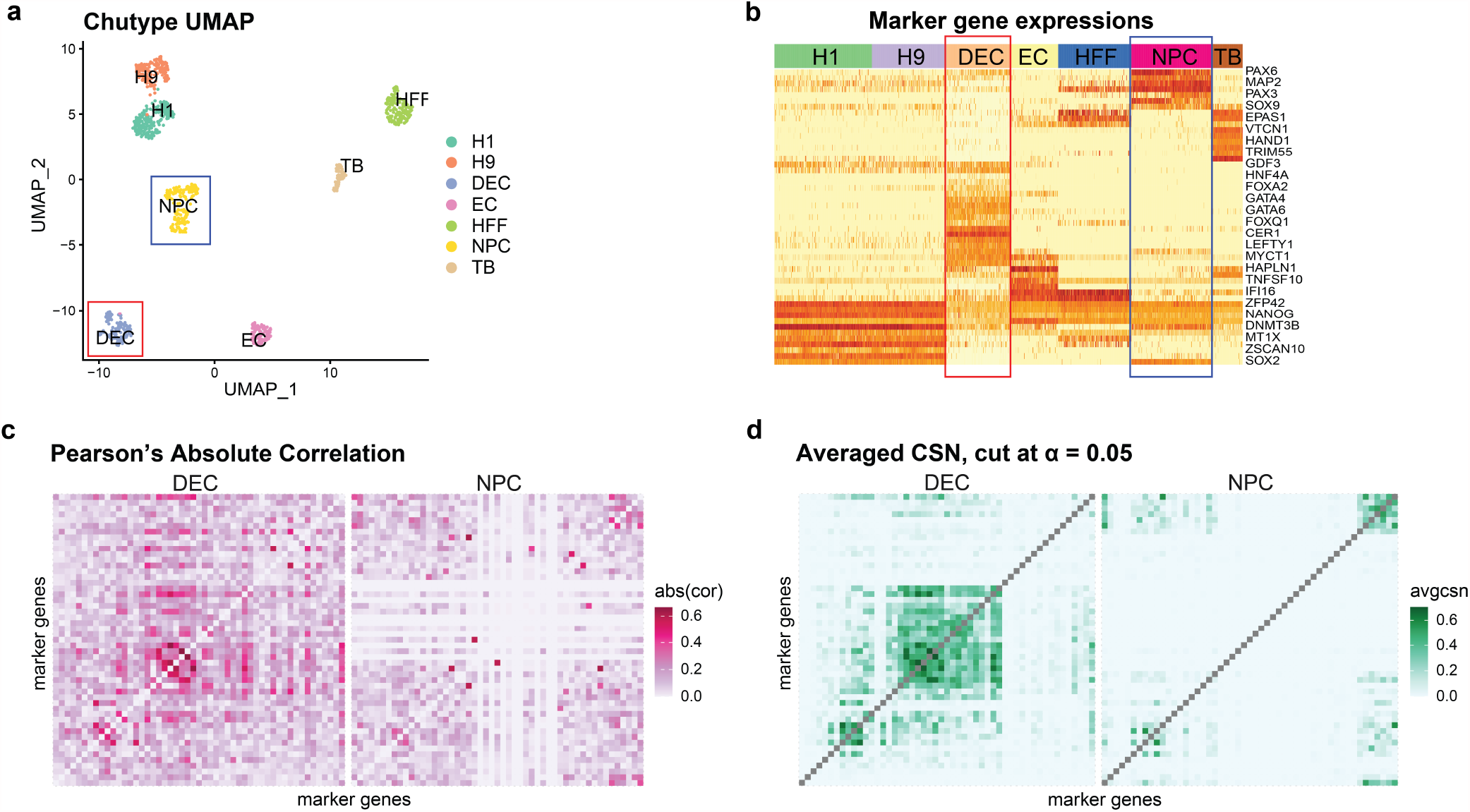
Estimated networks for Chutype cells. (**a**) UMAP of Chutype cells, colored by cell-types. The red and blue boxes indicate selected cell-type: DEC and NPC; (**b**) Heatmap of gene expression for 57 developmental genes for 7 cell-types. High to low expression corresponds to red to light yellow in heatmap. 7 cell-types are color coded by the band on the top of the heatmap. The red and blue boxes indicate DEC and NPC cell-types; Heatmaps of absolute Pearson’s correlation for DEC and NPC cell-type, calculated independently within cell-types. The order of genes are the same as (**b**); (**d**) Heatmaps of averaged locCSN for DEC and NPC cell-types, thresholded at Gaussian distribution *α* = 0.05 quantile. The order of genes are the same as (**b**).

### CSN analysis of developmental trajectories

To illustrate the application of CSN to developing cells we feature the developmental trajectory of excitatory neuron cells in the Developing Cortex Atlas dataset, which includes Radial Gila and Progenitors (P), Intermediate Progenitors (IP), Maturing Excitatory (ExN), Migrating Excitatory (ExM), Excitatory Upper-layer enriched (ExM-U), and Excitatory Deep layer (ExDp), with cell-type labels determined by the authors (27) Additional file 1: Supplementary Note 5 and Table S4). Using Slingshot (32), we estimate the developmental path consists of two trajectories, one ending in upper layer (U-curve) and the other in deep layer (D-curve) excitatory neurons (Figure 3**a** and **b**).

**Figure 3:**
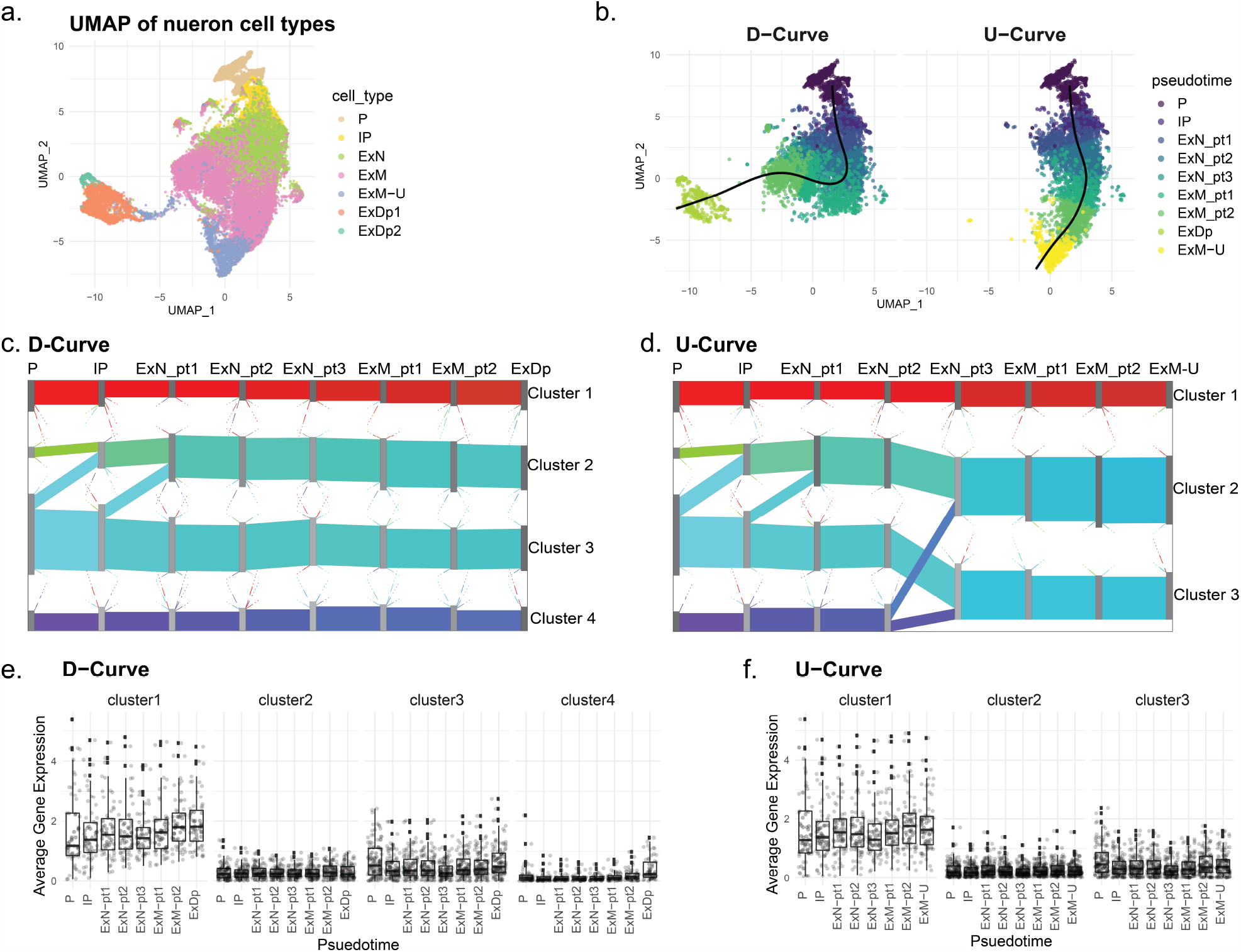
Development of networks in human fetal brain cells. (**a**) UMAP of human fetal brain single-cell expression from 7 cell-types involved in development of excitatory neuron cells, (**b**) with developmental trajectories superimposed. The UMAP plots indicate metacells, whose coordinates are the average UMAP cellular locations. The two principal curves generated by Slingshot (32). Colors are determined by the metacell’s pseudotime and cell types for the two curves, which are calculated as the average pseudotime over all cells in the metacell. The left and right panel shows the metacells assignment based on pseudotime to D-curve and U-curve, respectively. (**c**,**d**) Sankey plots of averaged CSN for 8 bins in D-curve and U-curve. Gene flows are shown as the colored band connecting two adjacent pseudotime bins for (**c**) D-curve and (**d**) U-curve. (**e**) and (**f**) depict boxplots of averaged metacell gene expression for 8 pseudotime bins in the final 4 and 3 clusters for the D- and U-curves, respectively. The x-axis shows pseudotime bins and y-axis shows the averaged expression.

**Figure 4:**
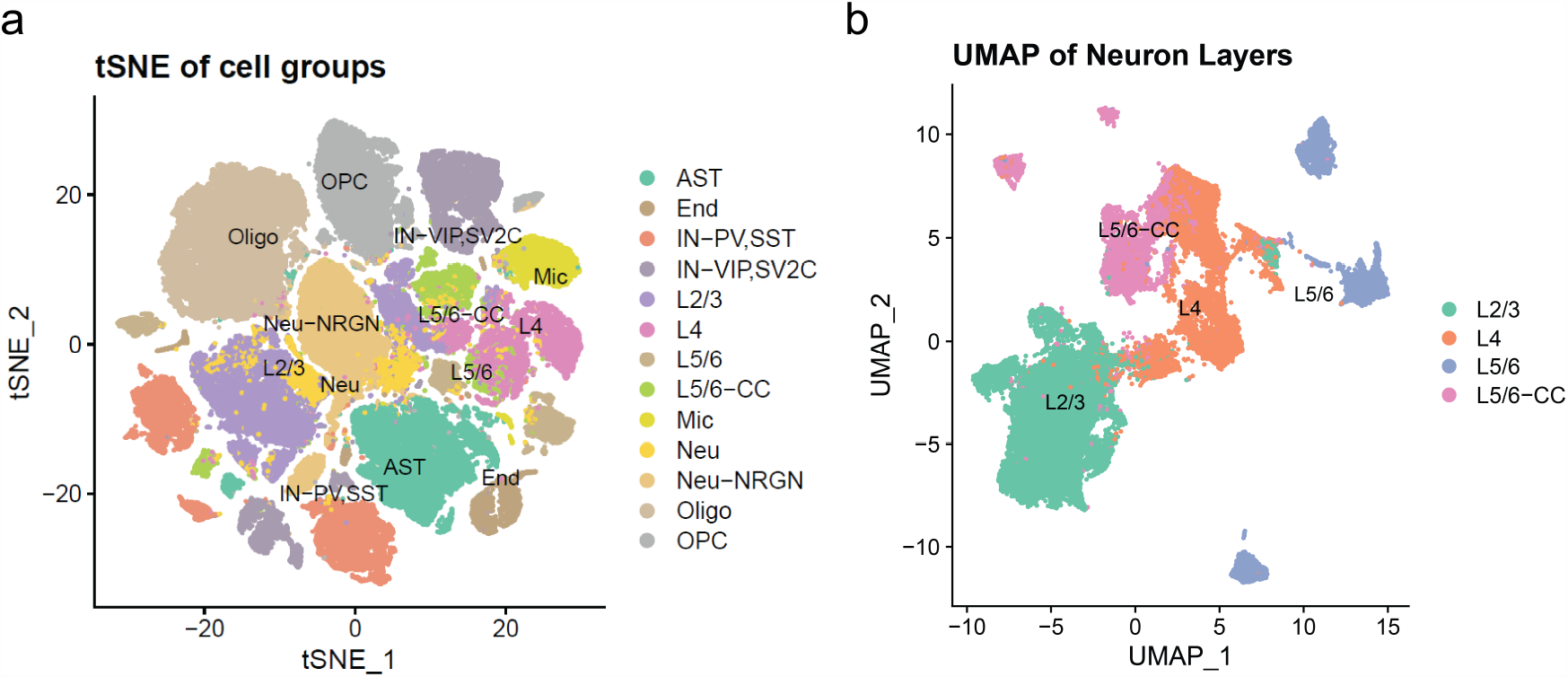
Dimension reduction of brain cells. (**a**) tSNE plot of all brain cells colored by cell-types. (**b**) UMAP plot of Neuron Layers(L) cells colored by 4 cell-types.

To circumvent challenges due to sparse counts, which are prevalent in these data, we pool similar cells within cell-type and form metacells (3), each containing approximately 20 cells. Each metacell’s expression, pseudotime and UMAP coordinate is computed as the average over pooled cells. Metacells are assigned to trajectories based on proximity using Slingshot (32). At the early stages of development the curves are nearly overlapping and metacells are not differentiated by curve, but as the cells develop, metacells can be assigned to a distinct curve (Additional file 1: Table S4-S6). Both curves start at the P cell class with full overlap, but moving along ExN and ExM cell classes, there is progressively less overlap between the two curves until they bifurcate, with D-curve and U-curve culminating in purely ExDp and ExM-U metacells, respectively. Next, within each cell-type, we generate bins based on the metacell pseudotime values. This effectively creates bins of approximately 800 metacells containing fairly homogeneous cells.

For each metacell, we use locCSN to compute gene networks. For illustration we focus on a restricted gene list. We choose ASD risk genes because the development of excitatory neurons is of interest in ASD research (27, 25, 31). Specifically CSNs were computed for 444 genes chosen by intersecting the expressed genes in the metacells with a list of 992 ASD-associated genes gleaned from the SFARI database (classes S, 1 and 2) (4). Our objective is to map the formation of gene-clusters over developmental epochs, as cells develop into upper or deep layer excitatory neurons. We apply PisCES (22) to the average CSNs per bin, to find time-varying gene-community structure in the D-curve and U-curve trajectories, and use Sankey plots to visualize the evolution of gene-communities (Additional file 1: Figure 3**c** and **d**). As expected the gene-communities are nearly identical for the two curves for the first four pseudotime bins, which share a large fraction of metacells, but the results progressively differentiate as the overlap in metacells diminishes. Most notably, the gene-community associated with cluster 4 (purple) for U-curve splits and merges into the other gene-communities at ExN pt3, while it persists for D-curve. For both curves, cluster 1 (red) contains genes that are more highly connected than the other loosely connected gene-communities (Additional file 1: Figure S10). For all of the remaining identified gene-clusters the correlation is extremely weak in the early stages and becomes more apparent in the final developmental bin.

We study the membership in gene-clusters when they are relatively stable without major splitting and merging from ExN pt3 an onward. We refer to gene-clusters in D-curve as cluster 1-4 and U-curve as cluster 1-3, where labels run from from top to bottom of the display. The numbers of genes in each stable cluster and the overlap between clusters from two curves are shown in the Table 1 (Additional file 2: Table S13). For example, most of the genes from the cluster 1 (dense cluster) for D-curve are included in the corresponding dense cluster, i.e. cluster 1 for U-curve. While the cluster membership is fairly stable along each curve, the strength of correlation changes over time (Additional file 1: Figure S10).

**Table 1:**
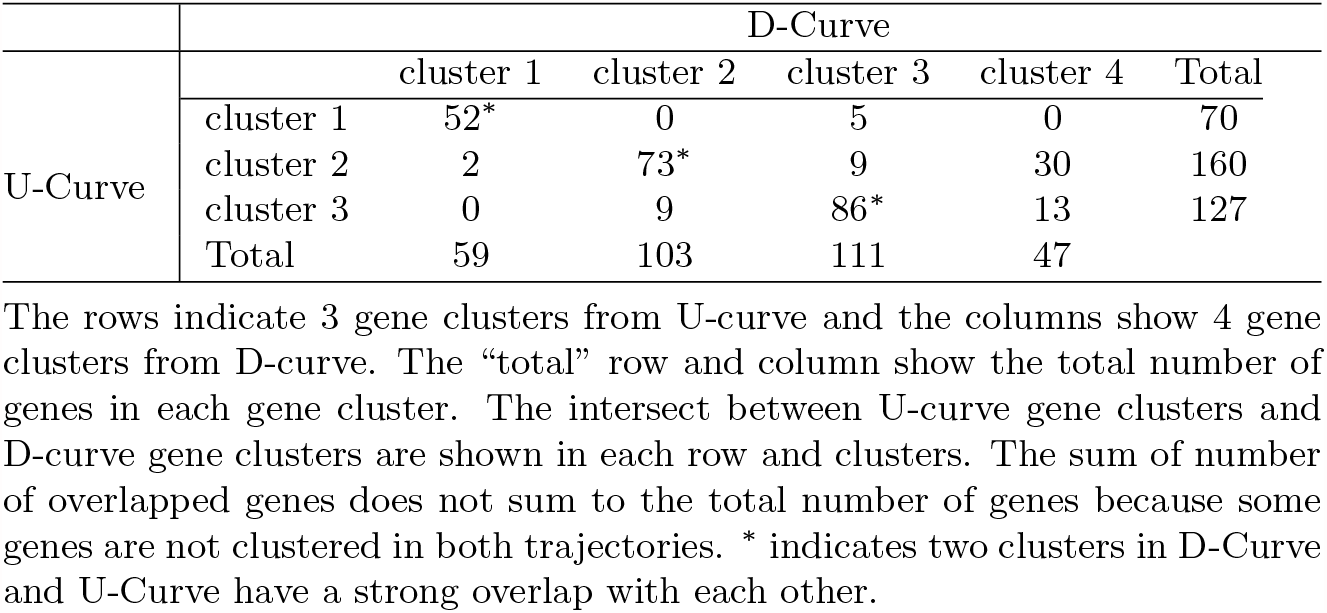
Number of genes in gene communities of D-curves and U-curve.

We also consider the averaged metacell gene expression across pseudotime for each gene-cluster (Figure 3**e** and **f**). Gene expression is relatively stable across cellular development. The dense gene-cluster (cluster 1) has high expressions while the loose gene-clusters have relatively low expression. Yet, even though all genes outside of the dense cluster have lower expression levels, using locCSN, combined with PisCES, we are able to detect subtle correlation and partition genes into gene-communities. It is worth noting that none of these communities are identified by WGCNA (19) (Additional file 1: Figure S11), which relies on Pearson’s correlation, but could also be implemented using average CSN matrices for potentially better performance.

To understand the function of gene communities, we check Gene Ontology (GO) terms (2) (Additional file 1: Supplementary Note 7) for the seven gene communities (Additional file 1: Figure S12, Additional file 3, Table S14). For both curves, GO term treemaps for the dense gene community (cluster 1) include metabolic processes, organelle organization and the process of mitosis, which are critical during the fetal stage. For D-curve, cluster 2 is enriched for chromatin organization, which is critical for gene expression regulation. While for U-curve, cluster 2 is enriched for organelle and cellular component organization. The most loosely structured set of genes, cluster 3 shows no GO enrichment for either curve suggesting it might not be biologically meaningful. Of greatest interest is cluster 4, which is ultimately restricted to D-curve and enriched for synaptic organization, suggesting that these neurons are more mature. This discovery fits with the biological process of neural migration which naturally proliferates the deep layers earlier than the upper layers.

### CSN analysis of brain cells from ASD subjects

To demonstrate how CSN is used to contrast network structure in two populations of cells, we analyze the ASD Brain dataset. These data feature single-nuclei RNA-seq (snRNA-seq) data from an ASD study of cortical nuclei assessing 13 cell-types and illustrate our results using excitatory neurons layers: L2/3, L4, L5/6 and L5/6-CC. For details of the full analysis pipeline and results see Additional file 1: Supplementary Note 6 and Figures S13. To illustrate the results, CSNs are computed only for genes implicated in ASD in the literature. We obtain the gene list by intersecting the 992 measured SFARI genes (4) with expressed genes of ASD case and control dataset and 942 genes remain.

After CSN construction, we compare the differences between control and ASD groups by testing for differences in their networks. We use two types of tests: first, an omnibus test for generic differences, and second, a targeted test, aimed at identifying high leverage genes that drive the difference. To test for differences in the distribution of the two classes, we can use a nonparametric distance-based test statistic (DISTp) taken from (23), coupled with a permutation test. This test is highly sensitive to differences in network structure, but a rejection merely indicates that the networks are significantly different. To gain further insights into the network differences, we can utilize the sparse-Leading-Eigenvalue-Driven (sLED) test (41). sLED is designed to detect signal attributable to a modest number of genes in the high dimensional setting encountered for studies of transcription. To emphasize the contrast with differentially expressed genes, we call these differential network genes. sLED takes as input a gene-gene relationship matrix such as the average CSN network or Pearson’s correlations matrix. (Further details can be found in Methods.)

The sLED-Pearson test detects no significant differences between control and ASD groups for any cell-type after adjusting for multiple testing, while the sLED-CSN and DISTp tests detect differences in 10 and 6 of the 13 cell-types tested, respectively (Table 2, Additional file 1: Table S10). For each cell-type that yielded a significant sLED-CSN difference we identify the high leverage genes (defined as the top loading genes that explain 90% of the signal by sLED, see Additional file 3: Table S15). We call these the “differential network (DN) genes”, to indicate that they strongly differ between case and control in their co-expression and network structure, but not necessarily in expression. The DN genes identified by sLED-CSN show striking differences in their averaged CSNs (Figure 5**a**-**d**, Additional file 1: Figure S15) and also their visualized networks (Figure 6). Removal of the DN genes reduces the number of detected differences from 10 to 2, suggesting that they explain much of the difference between the groups (Additional file 1: Table S11). Finally, comparing across layers, we find minimal overlap in the DN genes (Figure 7**a**) suggesting that the network differences are layer specific.

**Table 2:** P-values for comparing gene expressions and CSNs for control and ASD groups by cell-type.

| P-values | L2/3 | L4 | L5/6 | L5/6-CC |
| --- | --- | --- | --- | --- |
| sLED-CSN | 0.001* | 0.001* | 0.001* | 0.001* |
| sLED-Pearson | 0.089 | 0.861 | 0.564 | 0.348 |
| DISTp | 0.001* | 0.039 | 0.001* | 0.001* |

Comparing gene expressions and CSNs for control and ASD groups, p-values are calculated by sLED-CSN, sLED-Pearson and DISTp for 4 excitatory neuron layers: L2/3, L4, L5/6 and L5/6-CC. \* indicates significant differences between two groups after adjusting for multiple testing with 13 cell-types.

**Figure 5:**
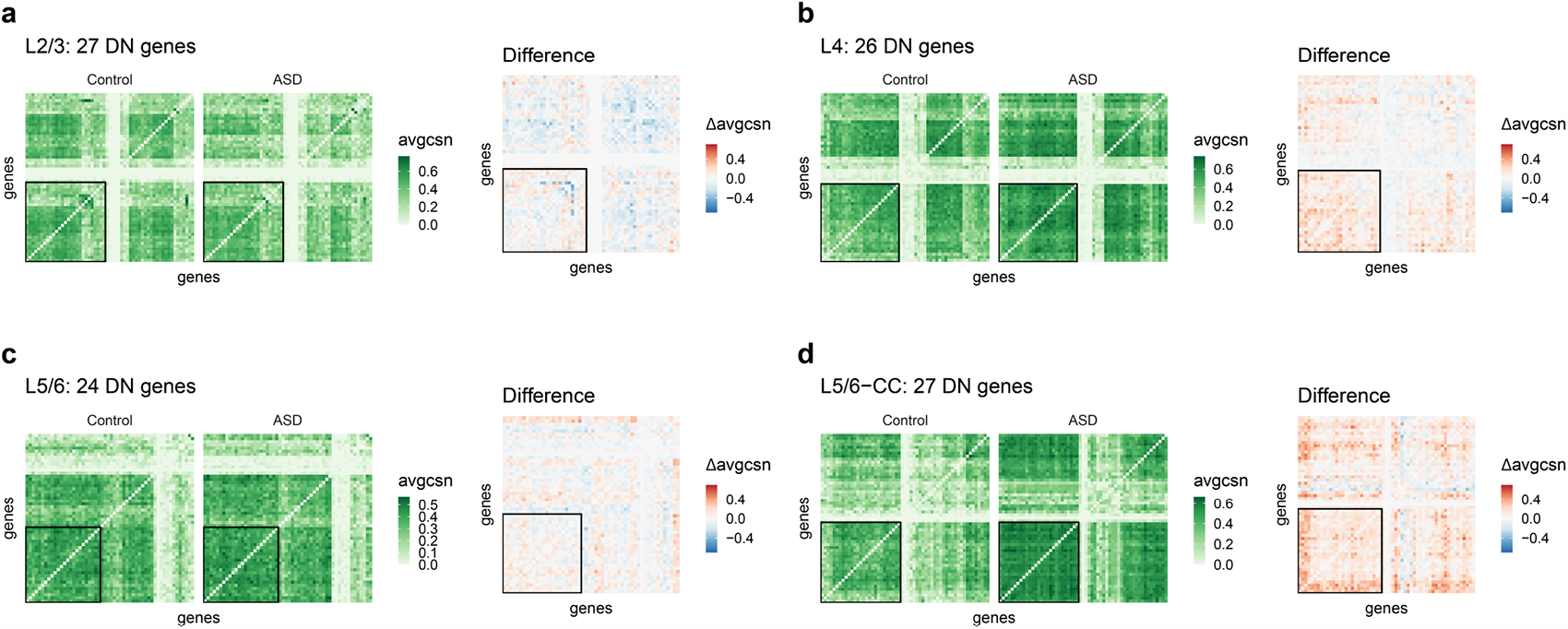
Heatmaps of average CSNs and the average difference (ASD minus Control). Heatmaps display sLED-CSN selected DN genes for each cell-type, contrasted with an additional 30 randomly selected genes from the 942 ASD genes. The black squares delineate the DN genes for each cell-types: (**a**) L2/3; (**b**) L4; (c) L5/6 ;and (**d**) L5/6-CC.

**Figure 6:**
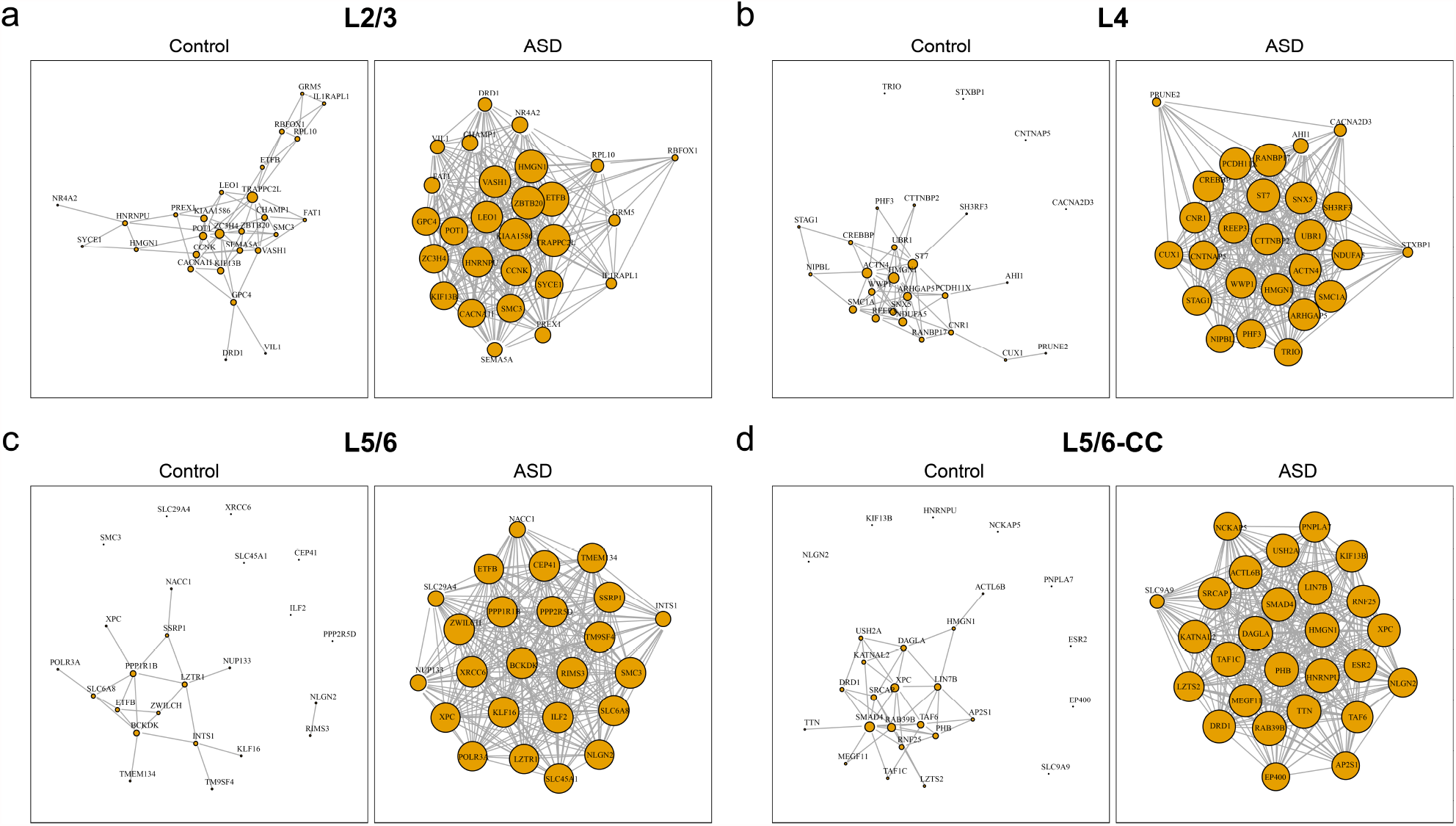
Gene networks for DN genes in the excitatory neuron layers. The networks are generated from averaged CSN of control and ASD groups. (**a**) L2/3; (**b**) L4; (**c**) L5/6; (**d**) L5/6-CC.

**Figure 7:**
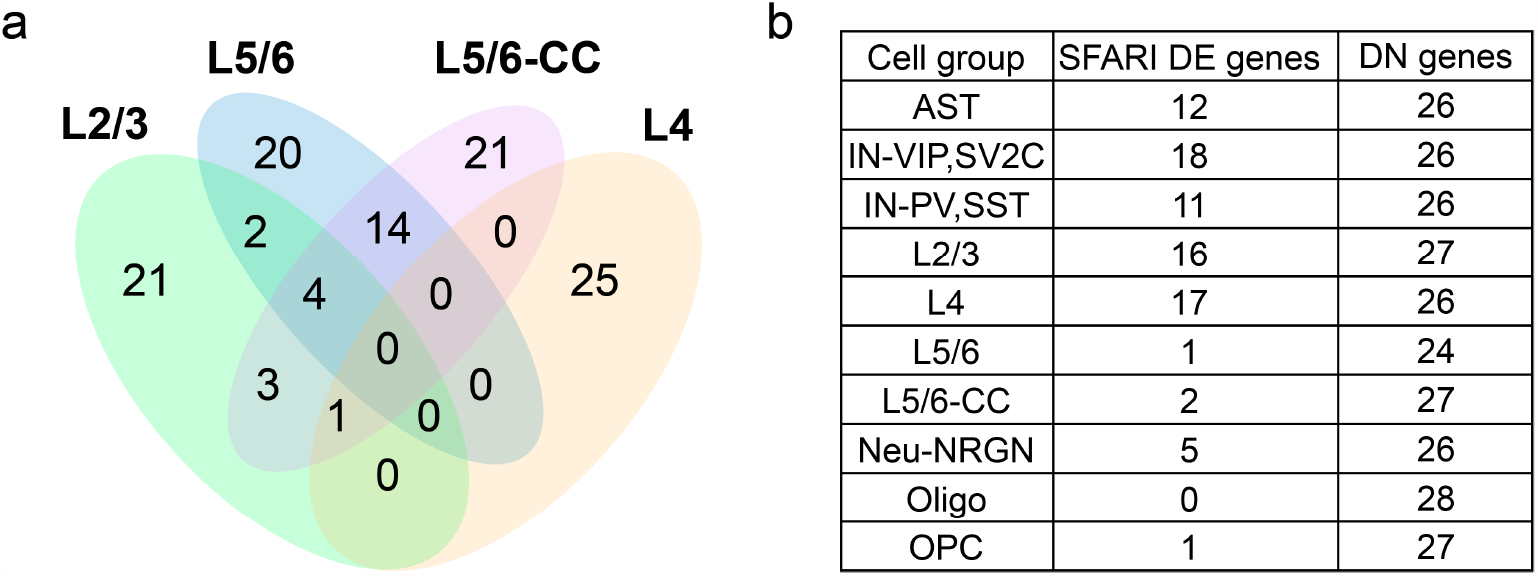
Contrasting differentially expressed and DN genes for ASD vs. Controls. (**a**) Venn diagram of all differential network (DN) genes for neuron layer cell-types. (**b**) Numbers of ASD differentially expressed (DE) genes and DN genes, which have no overlap.

For the 10 significant cell-types detected by sLED-CSN, we investigate the overlap between the differential expressed (DE) SFARI genes provided in (37) and the sLED-CSN DN genes. There is surprisingly no overlap (Figure 7**b**), suggesting that most DN genes do not show practically significant differences in expression level between control and ASD groups (Additional file 3: Table S15); similar conclusions can also be drawn by directly inspecting the marginal expression levels (Additional file 1: Figure S16). Overall the DN genes offer critical insights about transcription patterns and how they differ between ASD and control brains.

## Methods

### locCSN method

To compute 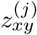, the estimated gene-gene relationship of cell *j* for gene pair (*x, y*), we first identify the neighborhood and then construct a test statistic. Let 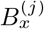 and 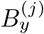 denote one-dimensional bins of cells for genes *x* and *y* centered at the expression levels for cell *j*, with widths *w*_*x*_ and *w*_*y*_:

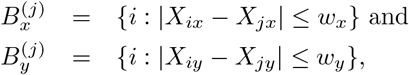

with cell counts 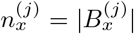 and 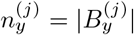. Let 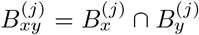 denote the joint window centered at cell *j* with counts 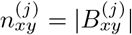.

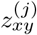 is represented by a normalized test statistic, 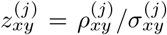, where 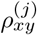 is a local test statistic for independence of genes *x* and *y*, comparing the joint distribution with the product of the marginals,

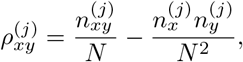

and the normalizing factor 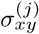 is the asymptotic standard deviation of 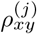 under the null hypothesis that genes *x* and *y* are independent, shown in (7) to equal

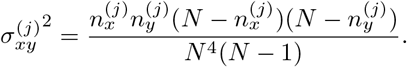

We remark that as a pre-processing step, we fix 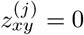 if the expression of either gene is zero for that cell.

The choice of the window sizes *w*_*x*_ and *w*_*y*_ plays an important role in the performance of the algorithm. Whereas oCSN by Dai et al. (7) uses window sizes equal to a fixed quantile range, we instead choose the window size based on a local standard deviation, which we compute iteratively. Bins 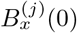 and 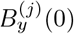 are initialized to include a quantile range (as in (7)), and then iteratively we follow

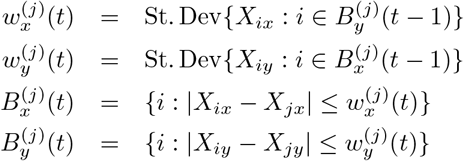

for *t* = 1, … until convergence is achieved (Additional file 1: Figure S2**b**). In practice, if the iterations do not converge, 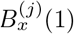 and 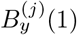 are used as window sizes.

### Metacell: Reduce sparsity of expressions

For single-cell and single-nuclei RNA-seq datasets, the expression can be very sparse. In this setting a direct application of the CSN algorithm fails to discover network structure. It can be advantageous to cluster the data before performing downstream analysis (18), hence, we apply the Metacell algorithm (3) before constructing CSNs. Metacell partitions cells into metacells, defined as disjoint clusters of homogeneous profiles. After applying Metacell to pre-labeled cells, we further divide metacells with multiple cell-types or subtypes into pure-cell-type metacells. Expression of a metacell is defined as the mean of the cells in the cluster, which alleviates the problem of having zero expression for many genes per cell. The metacells are then treated as cells for the purpose of constructing metacell-specific-networks. In this paper, for convenience, CSN refers to either a cell-specific-network or metacell-specific-network.

### Trajectory-based community detection

The PiSCES algorithm (22) is designed to identify cluster structure that varies smoothly over adjacent developmental periods. Following CSN construction for a smooth trajectory, where the cells have been binned by similar pseudotimes, we apply the PiSCES algorithm using the average CSN of each bin as input. The time-varying community structure can then be visualized using Sankey plots.

### Two sample testing

Given i.i.d. samples of expression levels, the computed CSNs for a set of cells are exchangeable, and hence permutation testing can be used to test for differences in CSN distribution. For this purpose, we suggest two types of tests: first, an omnibus test for generic differences, and second, a targeted test, aimed at identifying high leverage genes that drive the difference.

### DISTp: Test CSN differences between groups

Each cell’s adjacency matrix can be represented as a vector by converting matrices into vectors, resulting in *N*_1_ sample vectors from class 1 and *N*_2_ sample vectors from class 2. Let 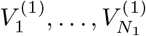 denote the vectorized adjacency matrices from class 1 and 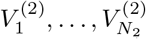 denote the same from class 2. The test statistic *Q* is a scaled *q*-norm divergence measurement, with *q* ∈ (0, 2) recommended (23), and is given by

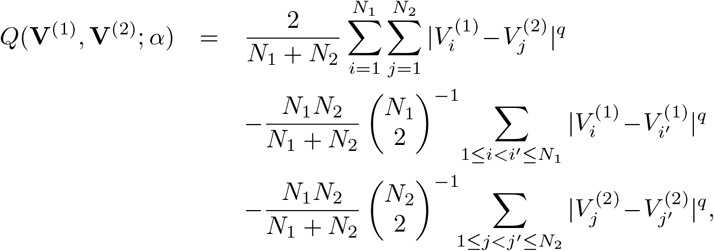

with p-value calculated by permutation test.

### sLED: Identify differential network genes

The sLED test relies on the same principles as Sparse Principal Component Analysis (SPCA), and was originally proposed for the difference in the Pearson’s correlation matrices of the two classes (sLED-Pearson). Here we instead propose using the difference in the average CSN as the test input (sLED-CSN). Given *N*_1_ CSN adjacency matrices from class 1 and *N*_2_ from class 2, denoted by 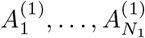 and 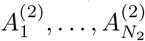, let *D* denote the difference between the average CSN for each class, so that 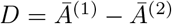. Then *D* can be used as the input to sLED, in which case the test statistic is computed from the spectrum of *D*. Additionally, the test also identifies a small cluster of leverage genes corresponding to the non-zero entries of the sparse leading eigenvector. The differential network genes are the ones that explain 90% of the variability among the leverage genes. These are candidate genes that have altered co-expression structure between the two groups. As with DISTp, the p-value of the test statistic is determined by permuting samples among cell classes.

A summary of notation that appears in Results and Method can be found in Additional file 1: Table S2.

### Datasets for analysis

#### Chu-type dataset

Chu et al.(6) includes 1018 cells and seven cell-types. This dataset contained the cells of human embryonic stem cell-derived lineage-specific progenitors. The cell-types including H1 embryonic stem cells, H9 embryonic stem cells, human foreskin fibroblasts (HFF), neuronal progenitor cells (NPC), definitive endoderm cells (DEC), endothelial cells (EC) and trophoblast-like cells (TB) were identified by fluorescence-activated cell sorting (FACS) with their respective markers. 9600 genes are obtained per cell on average.

#### Developing Cortex Atlas dataset

Polioudakis et al.(27) includes cells from mid-gestational human cortex. These data are derived from ∼40,000 cells from germinal zones (ventricular zone [VZ], subventricular zone [SVZ]), developing cortex (subplate [SP] and cortical plate [CP]) separated before single cell isolation.

#### Autism Spectrum Disorder (ASD) Brain dataset

Velmeshev et al.(37) includes snRNA-seq data from an ASD study, which collected 105 thousand nuclei from cortical samples taken from 22 ASD and 19 control samples from subjects between 4 and 22 years old. Samples were matched for age, sex, RNA integrity number, and postmortem interval.

### Runtime of locCSN

The CSN approach can be computationally intensive because we compute a test statistics for each pair of genes and each cell; however a number of speed ups are possible. By replacing cells with metacells (3), we can reduce the computational complexity substantially. We also provide an approximate CSN calculation by partitioning the outcome space for each pair of genes into a grid. Cells that fall into the same grid yield the same test statistic. With these approximations CSN can be readily applied to very large datasets with good accuracy (Additional file 1: Figure S17). Computational time can also be reduced by limiting the number of genes. While a large number of genes are expressed overall, a relatively modest number of genes are expressed in each cell type and it is natural to restrict analysis to this subset. Another option is to limit investigation to genes of particular biological interest, such as those implicated in disease or involved in biological processes of interest. Alternatively, to construct a network for a large number of genes (≈ 1000), the CSN computation can be applied to each gene pair in parallel.

We observe that, in concordance with the computational complexity, time consumption increases quadratically in the number of genes and linearly in the number of cells or metacell entries. Memory consumption increases in a similar manner. The examples of runtime against input matrix size are shown in Additional file 1: Table S12. Runtimes are measured under Python 3.7.6 [MSC v.1916 32 bit (Intel)].

## Discussion

Single-cell gene co-expression networks have the potential to yield critical insights into biological processes. Subtle differences in network structure can be used to classify cells into subtypes and by cell state. While each CSN is estimated with considerable noise, averaging CSNs over homogeneous cells can provide stable estimates of network structure and this can provide insights into how these networks vary by cell type, cell state and over developmental epochs.

Understanding cell-type specific gene networks can contribute substantially to our understanding of how biological processes are impacted by disease and disorders (24, 41, 13, 27). Just as we can test for differential gene expression, we can test for differential coexpression and aim to detect genes that drive the differences in coexpression. Small changes in gene expression can lead to substantial changes in network structure, ultimately with large biological effect. These differential network genes were called ‘dark’ genes by Dai et al. because we fail to detect them with traditional differential expression and yet can detect them with CSN analysis. For example, we found that most genes that leverage significant differences of connection between ASD and control groups are missed when simply comparing their expression levels. In a similar setting, using the scHOT algorithm, Ghazanfar et al.(11) identified DN genes in developing mouse liver cells that were not DE genes. Both analyses suggest that traditional analyses of gene expression miss critical signals about gene expression differences across developmental epochs and between phenotypes. CSN offers a powerful framework to discover DN genes.

CRISPR-Cas technologies (8, 9, 16) provide researchers with tools to introduce and assess the effects of many genetic perturbations. For example, (17) used the Perturb-Seq method (9) to introduce dozens of ASD risk genes to developing mice brains and then assessed the impact on the single-cell transcriptome. Because CSN estimates the network of each cell it is a natural tool for analysis of such perturbation experiments, which target individual cells with distinct perturbations. CSN analysis provides a useful tool for determining how such perturbations impact the network structure of a cell.

Constructing gene networks using the CSN approach is computationally intensive because we compute a test statistics for each pair of genes and each cell; however, a number of approximations can be readily applied to enhance the speed so that CSN can be applied to very large datasets. It is often natural to reduce the genes under investigation by CSN to a meaningful subset, such as genes previously implicated in genetic risk, genes mapped to critical pathways, or highly variable genes. Restricting the investigation to a subset of genes greatly reduces the computational complexity of CSN analysis, but more importantly, it can reveal more scientifically interpretable results. By focusing on hundreds of documented ASD risk genes, we were able to identify intriguing network structure in developmental trajectories and changes in network structure between ASD and control subjects. In the literature several papers have implicated particular cell-types, especially neurons and subtypes of neurons, in ASD risk (1, 26, 27, 31, 37, 38), but no consensus has been reached. Here we found that gene network structure differed subtly between the developmental trajectories of fetal brain cells for upper layer and deep layer excitatory neurons. Specifically, while both trajectories revealed clustering of genes involved in gene expression regulation, only the deep layer trajectory showed clustering of synaptic genes. Identifying differences in gene networks, both over developmental epochs and between phenotypes can shed light on the genetic etiology of human phenotypes.

## Conclusions

locCSN is a refined approach for generating co-expression networks at the level of the individual cell by sharing strength at the level of the gene pair. The output of locCSN can be used to uncover differences in co-expression between cells, and as examples we have shown how changes in co-expression may occur over a developmental trajectory, and also between case and control samples from autism spectrum disorder subjects. Our results highlight the importance of gene co-expression in addition to well-studied gene expression and provide a new perspective for gene modules detection and gene regulation in biological processes. We have made locCSN available at https://github.com/xuranw/locCSN.

## Data availability

All data used in this manuscript is publicly available. The accession numbers and links of third-party high-throughput sequencing data obtained from the GEO and EBI database were listed in Additional file 1, Tables S3, respectively.

## Software

All of the code used in these analyses is covered by the MIT License and is available from GitHub: https://github.com/xuranw/locCSN.

## Acknowledgements

The authors are grateful to Kevin Lin and Jinjin Tian for helpful comments.

## Authors’ contributions

This study was conceived of and led by K.R., D.C. and X.W. X.W. designed the model and estimation algorithm, implemented the locCSN software, and led the data analysis. K.R. and D.C. provided scientific insight on downstream analysis methods and data interpretation. X.W., K.R., and D.C. wrote the original draft. The authors read and approved the final manuscript.

## Additional Files

**Additional file 1 — Supplementary Information**

PDF file of supplementary information.

**Additional file 2 — Table S13**

Genes in each cluster of trajectory from Developing Cortex Atlas dataset.

**Additional file 3 — Table S14**

GO terms of the stable genes in each cluster of trajectory from Developing Cortex Atlas dataset.

**Additional file 4 — Table S15**

Table of leverage genes and top 90% differential network genes from ASD Brain dataset.

