## Additional file 1 for "Constructing local Cell Sepcific Networks from Single Cell Data"

#### Supplementary Information

Xuran Wang[1], David Choi[2] and Kathryn Roeder[1,3,*]

[1]Department of Statistics and Data Science, Carnegie Mellon University, Pittsburgh, PA, 15213, USA,
[2] Heinz College, Carnegie Mellon University, Pittsburgh, PA 15213, USA and
[3]Computational Biology Department, Carnegie Mellon University, Pittsburgh, PA, 15213, USA

### Supplementary Information

#### Supplementary Note 1 Performance under a correlated bivariate normal

Figure S1**a** shows the results of oCSN [4] and locCSN on data simulated from a correlated bivariate normal distribution,

$$\begin{pmatrix} X \\ Y \end{pmatrix} \sim N(\begin{pmatrix} 0 \\ 0 \end{pmatrix}, \begin{pmatrix} 1 & \rho \\ \rho & 1 \end{pmatrix})$$

where $\rho = 0.4$. For this seemingly canonical example of dependence between variables, oCSN rejects the independence hypothesis for 16% percent of the data points, while locCSN shows greater power, rejecting 55%. The lower power of oCSN in this example can be attributed to the choice of a fixed quantile range for the window used to estimate the marginal and joint densities. In particular, in areas of high density, where oCSN's power is lowest, the window becomes extremely small (Figure S1**b**). Similar patterns can be found in Figure S1**c**, which shows that locCSN chooses a constant window size for correlated Gaussian data, and in Figure S1d, which shows p-values computed by oCSN and locCSN for correlated normal data with $\rho$ ranging between 0.1 and 0.9.

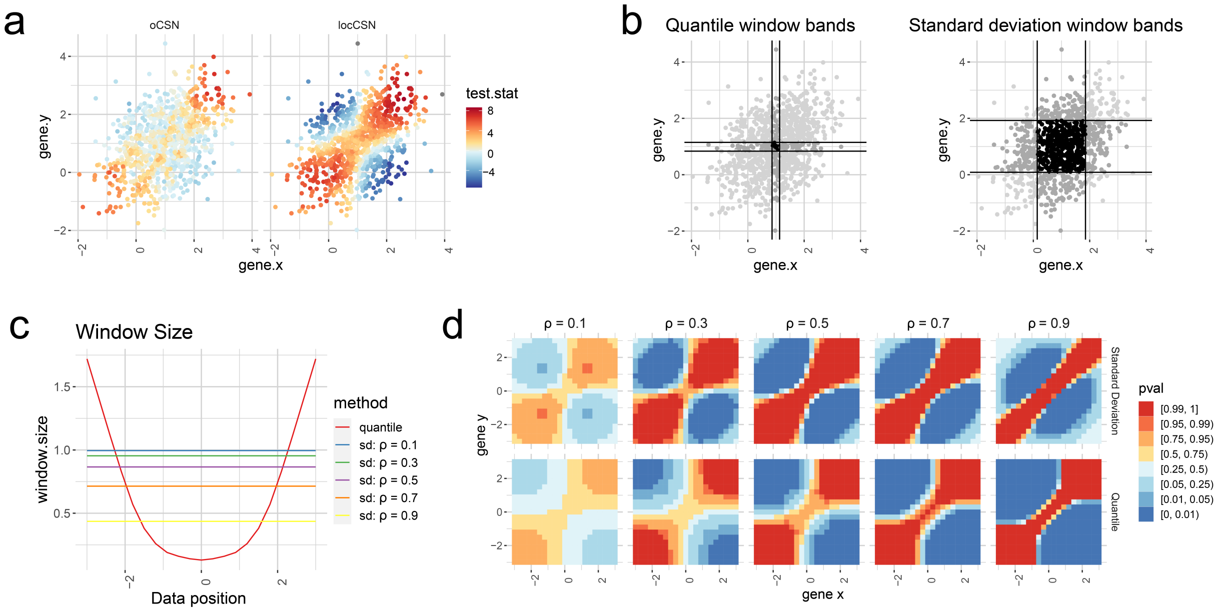


Figure S1: CSN window size analysis. (**a**) Scatter plot of dataset simulated from normal distribution with $\rho = 0.4$. The left panel is colored by the test statistics calculated from oCSN while the right panel is colored by the test statistics calculated from locCSN. (**b**) Scatter plot of the same dataset simulated from normal distribution. The left panel shows the 10% quantile window bands for a data point in the center. Dark gray shows points selected by gene $x$ band or gene $y$ band. Black shows points selected by both gene $x$ and gene $y$ bands. Light gray indicates points that are not selected by either bands. The right panel shows the scatter plot of the same dataset, window bands are derived from standard deviations. (**c**) Plots of Window sizes derived from quantile and standard deviation for normal distribution with different correlation $\rho$. The x-axis is the data point position and the y-axis is the window size. Different color shows window sizes derived from quantile or standard deviation under different correlations.

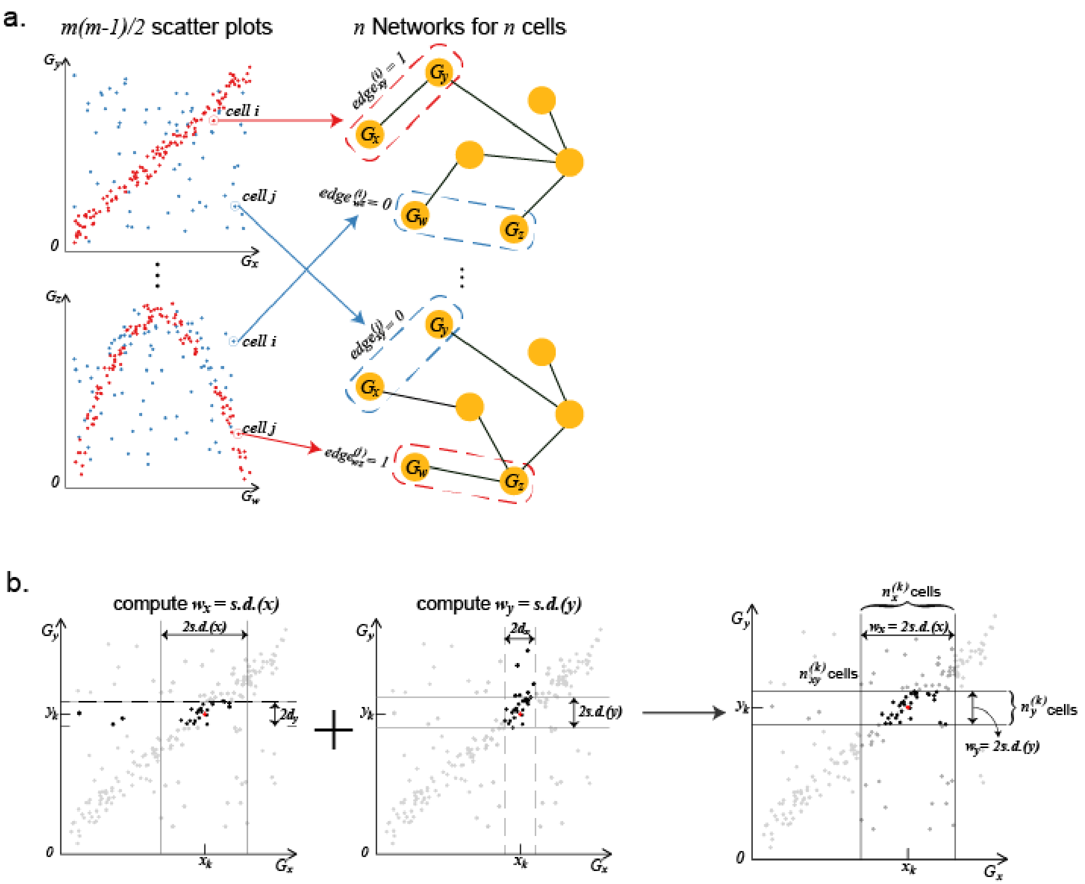


Figure S2: (**a**) Adapted from Dai et al.[4, Figure 1A]. Scatterplots of gene expression levels for gene pairs $(x, y)$ and $(w, z)$, showing regions of high and low density (red and blue) compared to the product of the marginal densities, with corresponding CSN edges or non-edges for these gene pairs highlighted for two cells $i$ and $j$. The connection between gene pairs are different across cells. (**b**) Standard deviation derived window size for one cell. The first scatter plot shows the quantile derived window for gene $y$, $w_y = 2d_y$. With this window size, we calculate the standard deviation of gene $x$ expression, using cells within this window size. We take the obtained standard deviation for gene $x$ as the window size for gene $x$, $w_x = 2s.d.(x)$. Second scatter plot shows the same thing as the first one with swapped $x$ and $y$. Finally, we use the window size derived from standard deviation for further calculation, that is $w_x = 2s.d.(x)$ and $w_y = 2s.d.(y)$.

#### Supplementary Note 2 Simulations study of CSN parameters

The choice of parameters influence the CSN performance. Using different connection strength and expression levels between genes, we study how different window size and threshold affect CSN performance. We simulated datasets from ESCO [8] with two different settings: 1. True counts without technical noise; 2. Counts with technical noise.

##### True counts without technical noise

Simulation results are generated from ESCO with a single cell group. There are 200 cells and 100 genes. The code to reproduce this simulation is here: code. The gene-gene correlation matrix exhibits block structure of the form (Figure S3**a**), or equivalently has off-diagonal entries for the 6 blocks given by the matrix

$$\begin{bmatrix} 0.9 & 0.7 & 0.5 & 0.3 & 0.1 & 0 \\ 0.7 & 0.9 & 0.7 & 0.5 & 0.3 & 0 \\ 0.5 & 0.7 & 0.9 & 0.7 & 0.5 & 0 \\ 0.3 & 0.5 & 0.7 & 0.9 & 0.7 & 0 \\ 0.1 & 0.3 & 0.5 & 0.7 & 0.9 & 0 \\ 0 & 0 & 0 & 0 & 0 & 0 \end{bmatrix},$$

where the first 5 blocks have 15 genes each, and the 6'th block has 25 genes that are independent from all others.

CSN are performed on log-transformed CPM with ESCO simulated read counts data. Figure S3**c** and **d** shows histograms of the CSN test statistic for gene pairs as a function of their correlation and the choice of window size, which was either a fixed quantile range (as suggested by [4]) of width 5%, 10%, 15%, or 20%; or else initialized to a fixed quantile range and then adapted using locCSN (resulting in "standard deviation window sizes"). We see that the test statistic is shifted for gene pairs that are highly correlated.

For the same choice of window sizes, Figure S4**a - c** show ROC and AOC for the task of discriminating gene pairs that are uncorrelated from those whose correlation is $\geq 0.5$. 2100 genes pairs were randomly selected for this task, balanced between the two categories. The curves show that for this task, the standard deviation-based window sizes used by locCSN perform better than the fixed quantile range windows proposed by [4] Figures S4**g** and **h** shows that the average CSN heatmaps using the standard deviation-based window size resemble the original connection matrix (Figure S3**a**) more strongly that do those using the quantile-based window sizes.

##### Counts with technical noise

To better represent single-cell dataset, which are extremely sparse, we use the down-sampling feature in ESCO to produce realistic simulated data. Increasing sparsity weakens the strength of connection in the observed datasets, so we use correlation matrices with stronger values to produce meaningful results. Simulation is again performed by ESCO with 200 cells and 100 genes. We study 2 scenarios:

1. Strong connection: there are 4 blocks of genes, 25 genes each. Within each block, genes are highly correlated with $\rho = 0.95$. Genes from different blocks are independent (Figure S5**a**);

2. With weaker connection: same as above, but blocks 3 and 4 are not independent, and instead have a weaker correlation $\rho = 0.5$ (Figure S6**a**).

The parameters for ESCO simulation with down-sampling are set at lib.loc = 7 and alpha_mean = 0.7. lib.loc indicates the overall expression level of the datasets and alpha_mean controls the strength of down-sampling. 40%-50% of the simulated expression are zeros, which approximately corresponds to single-cell RNA-sequencing data with high depth for a particular cell-type. The code to reproduce two simulation scenarios is here: code.

Figures S5**c-e** and S6**c-e** show ROC curves and ACC for the task of discriminating gene pairs that are uncorrelated from those with positive correlation. Results show better performance when using locCSN compared to using fixed quantile window sizes. We also show the heatmap of the average CSN (Figures S5**f** and S6**f**), thresholded at $\alpha = 0.01$ for strong connection scenario and at $\alpha = 0.05$ for the weaker connection scenario.

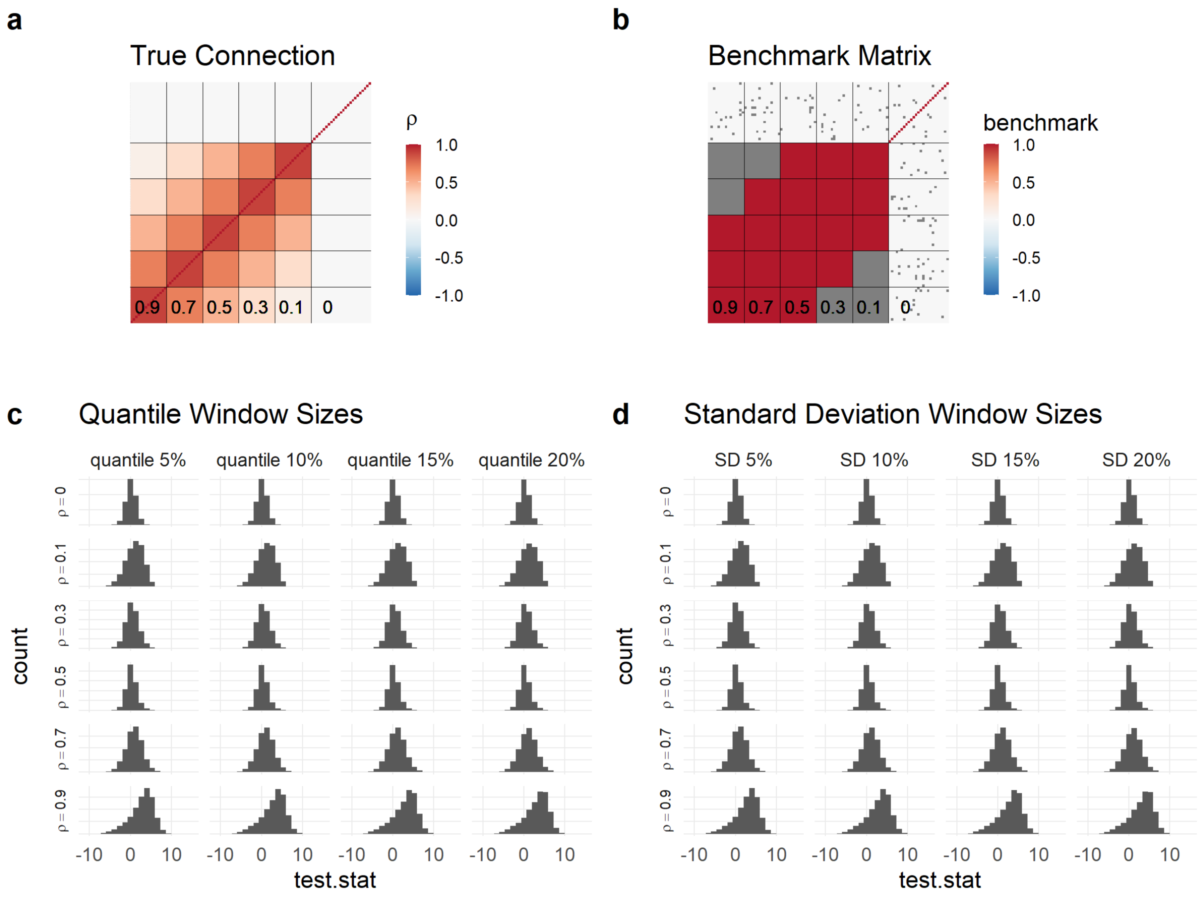


Figure S3: ESCO simulated single cell expressions without technical noise. (**a**) Heatmap of block structure in gene-gene correlation matrix used for ESCO simulation. (**b**) Gene pairs used for task of classifying pairs with correlation $\rho \geq 0.5$ vs $\rho = 0$. (**c** and **d**) Histogram of CSN test statistics. We exclude genes that are not expressed for all cells. The x-axis shows the CSN test statistics and y -axis is the counts of test statistics. We separate different correlations $\rho$ with rows and different window sizes are shown in columns. (**c**) Histogram for quantile window sizes, 5%, 10%, 15% and 20%. (**d**) Histogram for standard deviation (SD) with starting window size at 5%, 10%, 15% and 20%.

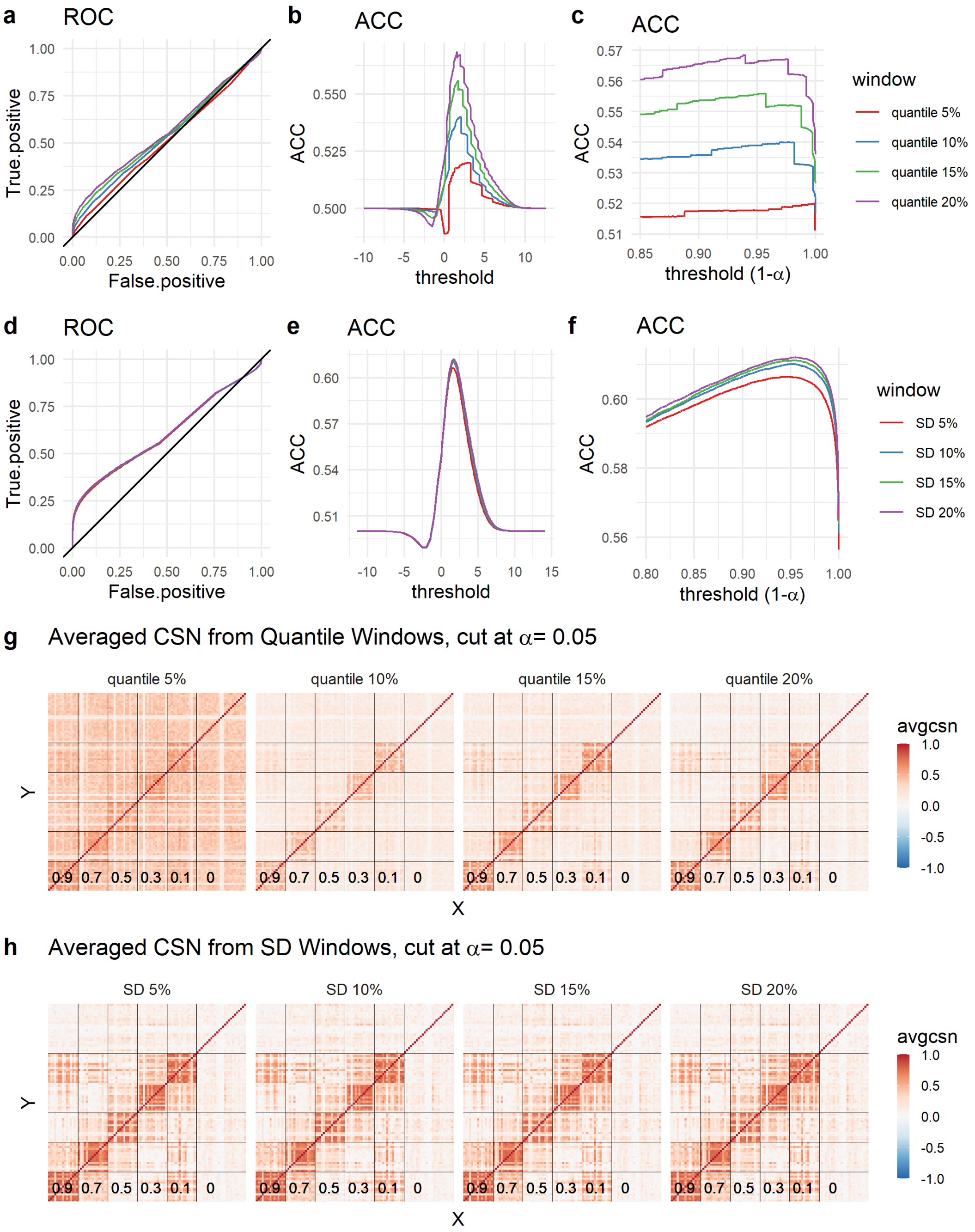


Figure S4: Evaluation of CSN test statistics from simulated dataset. (**a - c**) Evaluations of quantile window sizes. (**a**) ROC curve; (**b**) Accuracy (ACC) curve; (**c**) ACC curve with threshold $Z_{(1-\alpha)}$. The x-axis is $1-\alpha$. (**d - f**) Evaluations of quantile window sizes. (**d**) ROC curve; (**e**) ACC curve; (**f**) ACC curve with threshold $Z_{(1-\alpha)}$. The x-axis is $1-\alpha$. (**g - h**) Heatmaps of averaged CSN. Each panel shows averaged CSN for a specific window size. The threshold for having edges is $\alpha = 0.05$. (**g**) Heatmaps of averaged CSN for quantile windows; (**h**) Heatmaps of averaged CSN for standard deviation windows.

The two scenarios both support that standard deviation windows work better than quantile windows in terms of false discovery and accuracy. The simulation also suggests that the choice of threshold also depends on how true connections are defined. If we only consider strong connections (correlation $> 0.9$) as connected, we can use larger threshold. On the other hand, if we want to include medium strength connections (correlation $\geq 0.5$) as connected, we can use the smaller threshold.

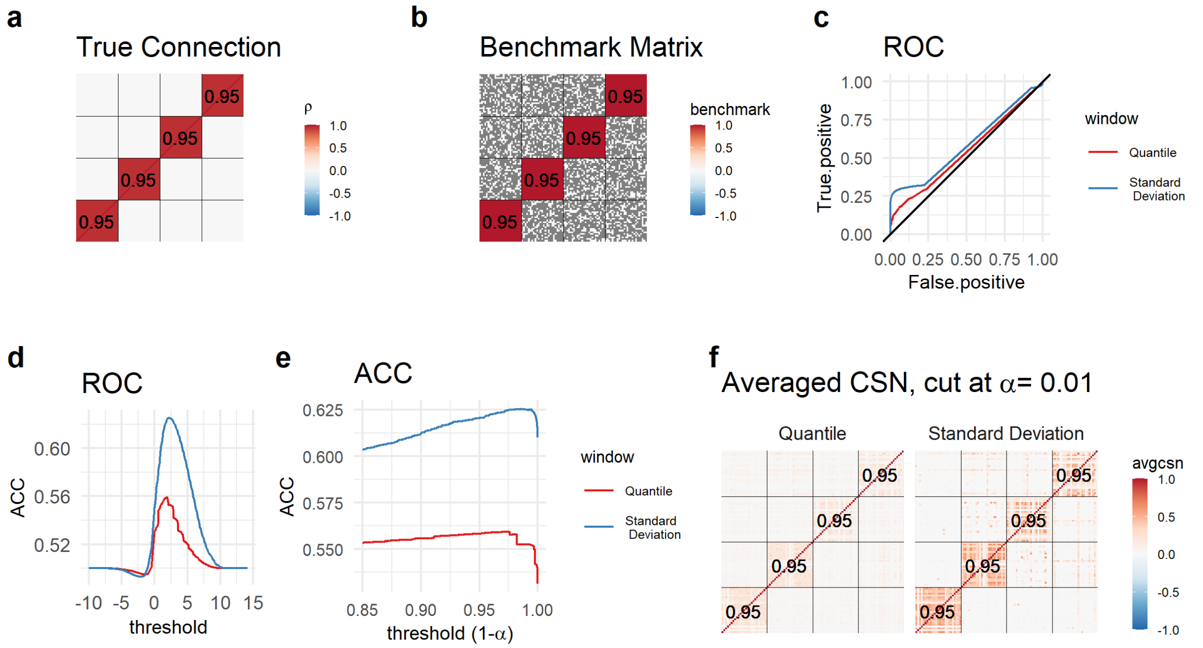


Figure S5: Evaluation of ESCO simulated dataset with strong connections and technical noise. (**a**) Heatmap of true correlation matrix; (**b**) Gene pairs used for classification task; (**c** - **e**) Evaluation curves for quantile and standard deviation window sizes. (**c**) ROC curve; (**d**) ACC curve; (**e**) ACC curve with threshold $Z_{(1-\alpha)}$. The x-axis is $1 - \alpha$. (**f**) Heatmaps of averaged CSN with threshold at $\alpha = 0.01$. Two panels indicate quantile window and standard deviation window.

##### Comparison between Pearson's correlation and CSNs

Using the same simulation with technical noise as above, we compare the empirical Pearson's correlation matrix and the averaged CSNs to the true block-structured correlation matrix. CSNs are calculated using standard deviation window size and averaged CSNs are thresholded at $\alpha = 0.01$ and $\alpha = 0.05$. Figure S7 shows the heatmaps of true and estimated matrices for two scenarios.

Now we use $L_1$-norm to measure the differences between the true correlation structure matrix ($A$) and the estimated version $\hat{A}$, which may be either the average CSN or the empirical Pearson's correlation matrix. In both scenarios, averaged CSN estimates $A$ better than the empirical Pearson's correlation, as can be seen in Table S1, and also visually in the heatmaps themselves.

##### Compare CSN with BigSCale correlation

Next we simulate data from ESCO, with 10000 genes and 2500 cells. We focus on 125 housekeeping genes that are not correlated with each other. After down-sampling the read counts to weaken the signal, we compare BigSCale [5] and locCSN using metacells. With no correlation between genes, we should not detect connections between genes. From 2500 cells, we constructed 158 metacells and on average, there are 15 cells per metacell. The heatmaps of

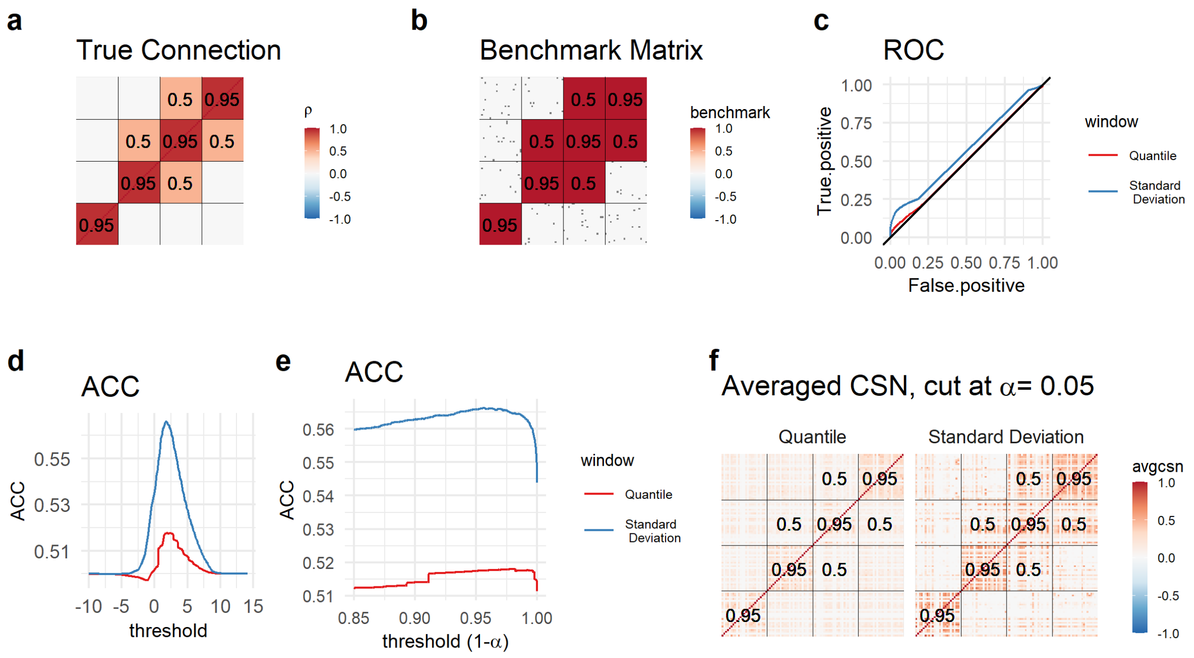


Figure S6: Evaluation of ESCO simulated dataset with weaker connections and technical noise. (**a**) Heatmap of true correlation matrix; (**b**) Gene pairs used for classification task; (**c** - **e**) Evaluation curves for quantile and standard deviation window sizes. (**c**) ROC curve; (**d**) ACC curve; (**e**) ACC curve with threshold $Z_{(1-\alpha)}$. The x-axis is $1-\alpha$. (**f**) Heatmaps of averaged CSN with threshold at $\alpha = 0.05$. Two panels indicate quantile window and standard deviation window.

| | Strong Connection | With Weaker Connection |
|---|---|---|
| Average CSN ($\alpha = 0.05$) | 30.73 | 46.73 |
| Average CSN ($\alpha = 0.01$) | 27.67 | 46.95 |
| Pearson's Correlation | 37.66 | 50.42 |

Table S1: Distances between true connection matrix and estimated matrices, measured by $L_1$ norm.

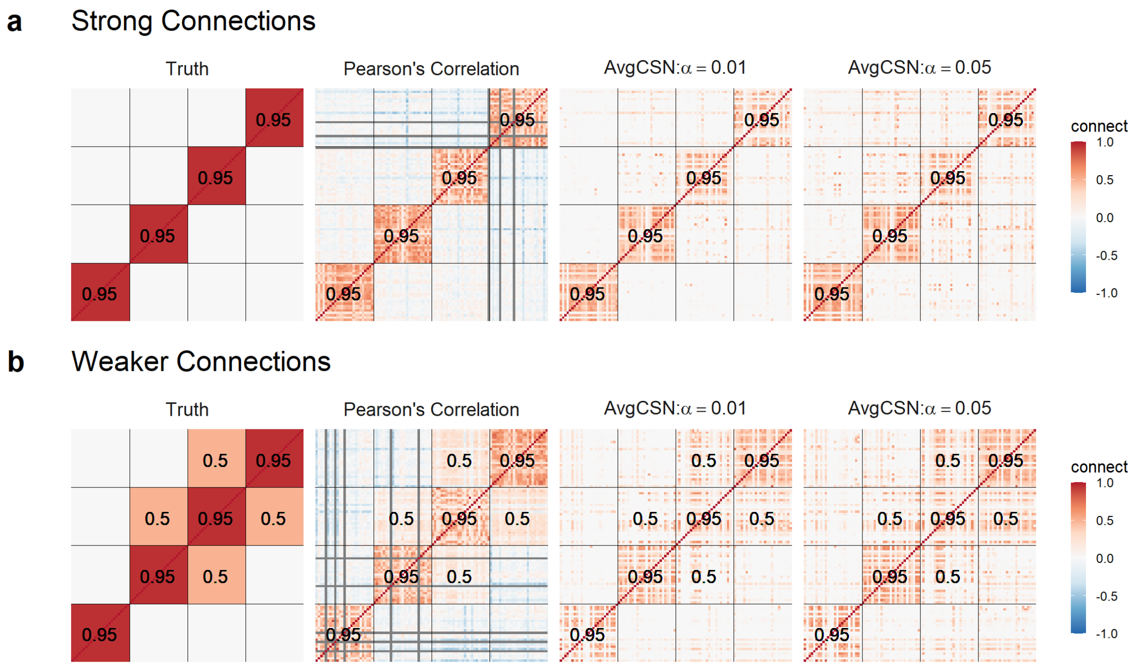


Figure S7: True correlations and estimates using either empirical correlation or average CSN. The first panel shows true correlations and the following 3 panels shows the estimates. The second panel shows empirical Pearson's correlation. The next two panels are the averaged CSN with threshold $\alpha = 0.01$ and $\alpha = 0.05$. (**a**) Strong connections; (**b**) Weaker connections.

Pearson's correlation of down-sampled and true read counts are shown in the first panel of Figure S8**f**. The BigSCale correlation shows false positives between genes when there are no connection between genes. By contrast, average CSN with metacells shows no connection between genes.

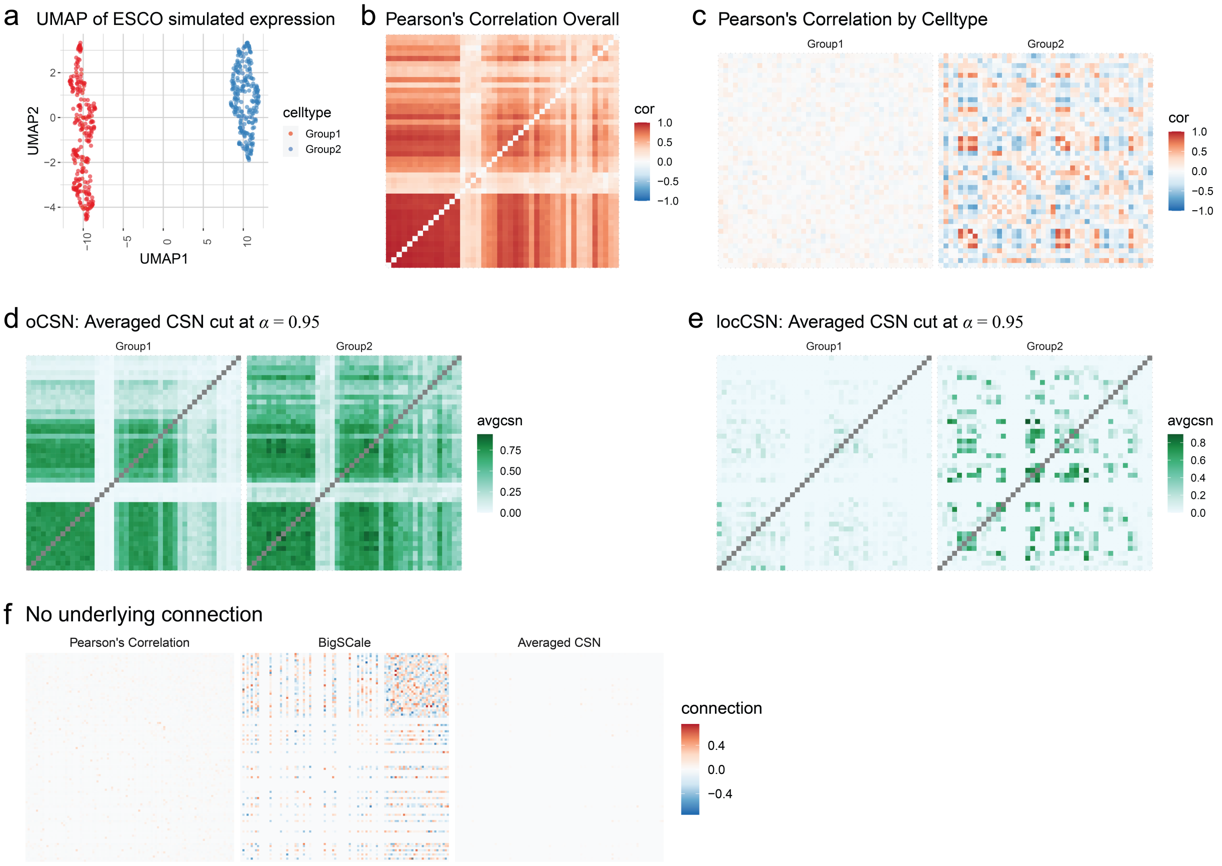


Figure S8: Network estimation for two simulated cell-types. (**a**) UMAP from ESCO simulated gene expression. (**b**) Heatmap of Pearson's correlations of genes, calculated ignoring cell-types. (**c**) Heatmaps of Pearson's correlations of genes, calculated independently for each cell-type. (d-e) Heatmaps of averaged CSN within cell-type, thresholded by $\alpha = 0.95$ quantile of standard normal distribution. (**d**) oCSN calculated ignoring cell-types. (**e**) locCSN calculated independently for each cell-type. (**f**) For a dataset simulated using ESCO with no correlation between any genes, heatmaps of Pearson's correlation, estimated BigSCale network and averaged locCSN.

#### Supplementary Note 3 Notations

| **Variable** | **Definition** |
|---|---|
| $N$, $N_1$ and $N_2$ | number of cells (total, class 1 and class 2). |
| $i$, $j$ | index of cell: cell $i$ and cell $j$. |
| $G$ | number of genes. |
| $x$, $y$ | index of gene: gene $x$ and gene $y$. |
| $w_x$, $w_y$ | width of window for gene $x$ and $y$. |
| $X_{jx}$ | gene expression of cell $j$ and gene $x$. |
| $B_x^{(j)}$, $B_y^{(j)}$, $B_{xy}^{(j)}$ | one-dimensional bins for gene $x$ at the expression level for cell $j$, with window $w_x$; for gene $y$ at expression level for cell $j$ with window $w_y$; $B_{xy}^{(j)}$ is the joint window. |
| $n_x^{(j)}$, $n_y^{(j)}$ and $n_{xy}^{(j)}$ | number of cells in bins $B_x^{(j)}$, $B_y^{(j)}$ and $B_{xy}^{(j)}$. |
| $\rho_{xy}^{(j)}$ | local test statistics for independence of genes $x$ and $y$. |
| ${\sigma_{xy}^{(j)}}^2$ | asymptotic standard deviation. |
| $t$ | iteration in standard deviation window size calculation. |
| $z_{xy}^{(j)} = \rho_{xy}^{(j)}/\sigma_{xy}^{(j)}$ | normalized test statistics.for gene pair $(x, y)$ and cell $j$. |
| $A_j$, $A_j^{(1)}$ | Estimated adjacency matrix for cell $j$ and for cell $j$ in class 1. |
| $a_{xy}^{(j)}$ | Entry of $A_j$, gene pair $(x, y)$ and cell $j$. $a_{xy}^{(j)} = 0$ or 1. |
| $\alpha$ | Standard normal tail quantile. |
| $Z_{(1-\alpha)}$ | reverse CDF of standard normal at $1 - \alpha$. |
| $V^{(1)}$, $V_i^{(1)}$ | vectorized adjacency matrix for class 1 and for cell $i$ in class 1. |
| $q$ | $q$-norm for DIST-p distance. |
| $D$ | the differences between average CSN for each class, $D = \bar{A}^{(1)} - \bar{A}^{(2)}$. |

Table S2: Notations and Definitions

#### Supplementary Note 4 Data summary

| Datasets | ESCO Synthetic | Chutype | Brain Cortex Atlas | ASD Brain |
|---|---|---|---|---|
| References | Tian et al.(2020) [8] | Chu et al.(2016) [3] | Polioudakis et al.(2019) [6] | Velmeshev et al.(2019) [9] |
| Tissue | NA | Human Embryonic Stem Cells | Human fetal Brain Cortex | human brain |
| # cell | 2000 | 1018 | 25,013 | 104,559 |
| # cell-types | 2 | 7 | 16 | 18 |
| # genes | 100 | 16,619 | 35,543 | 41,202 |
| # genes for analysis | 30 markers | 51 developmental markers | 444 expressed SFARI ASD genes | 942 expressed SFARI ASD genes |
| Data Availability | Github* | GSE75748 | Website | PRJNA434002 |

Table S3: Data summary of single cell data for analysis. * The code to reproduce this dataset is here: code.

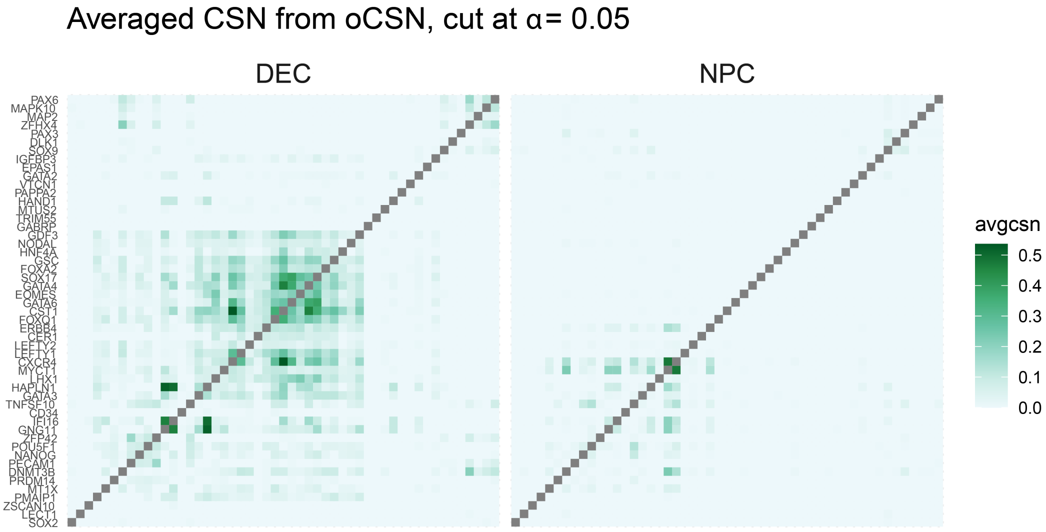


Figure S9: Heatmap of averaged CSN calculated from oCSN. Dataset is the same as in Figure 2, that is, Chutype DEC and NPC cell-type. The cut-off is $\alpha = 0.05$.

#### Supplementary Note 5 CSN trajectory analysis of Brain Cortex Atlas data: data processing

The human brain cortex atlas data contains 35,543 genes and 33,986 cells from 4 samples. There are 16 cell-types and each cell-type contains multiple subtypes. We focus on 10 cell types for analysis, specifically the 6 neuron cell types: ExDp1, ExDp2, ExM, ExM-U, ExN and IP, plus 4 radial glia and progenitors (P) cell-types: vRG, oRG, PgS and PgG2M. Metacells are constructed sample by sample and within a subtype. Based on Figure 3 ExDp1 and ExDp2 cell-types are combined as ExDp and the 4 radial glia and progenitors cell-types as P for further analysis (ExDp are subsequently partitioned at a later stage of analysis). Table S4 shows the number of cells and metacells for 7 major cell-types: P, IP, ExN, ExM, ExM-U and ExDp.

Prior to CSN construction along the curve, we generate metacell bins based on pseudotime of the curve within each cell-type. Each bin contains around 800 metacells, which are relatively homogeneous; however, for each cell-type, some metacells are deemed outliers based on their pseudotime scores. We retain metacells whose pseudotime are within 2 standard deviations of the mean within the bin. These are the cells that will be utilized for CSN construction. The number of bins used for each cell-type, and number of metacells (before and after outlier screening) are listed in Table S5. Since there are only 58 metacells for ExM-U in the leftmost curve, we ignore this cell-type for the D-Curve analysis. This cell-type properly belongs in the U-curve analysis. Table S6 shows the overlapping of metacells for the two curves. P metacells are shared across both curves. IP and ExN metacells are largely shared between the two curves. The split in trajectories occurs during development of the ExM cell-type, which show considerably less overlap of metacells between the two curves.

| Cell-type | P | IP | ExN | ExM | ExM-U | ExDp |
|---|---|---|---|---|---|---|
| Number of cells | 4204 | 2150 | 9995 | 9822 | 1756 | 2205 |
| Number of metacells | 720 | 574 | 2759 | 2415 | 424 | 271 |

Table S4: Number of cells and metacells in cell-types.

| Curves | D-Curve | | | | | | U-Curve | | | | |
|---|---|---|---|---|---|---|---|---|---|---|---|
| Cell type | P | IP | ExN | ExM | ExM-U | ExDp | P | IP | ExN | ExM | ExM-U |
| Number of metacells | 720 | 574 | 2451 | 1604 | 58 | 265 | 720 | 569 | 1913 | 1189 | 335 |
| Remaining metacells | 720 | 559 | 2373 | 1488 | 0 | 262 | 720 | 531 | 1804 | 1097 | 324 |
| Number of bins | 1 | 1 | 3 | 2 | 0 | 1 | 1 | 1 | 3 | 2 | 1 |

Table S5: Number of metacells in two curves for 6 cell-types. Before and after removal of pseudotime outliers.

| Number of metacells | P | IP | ExN | ExM |
|---|---|---|---|---|
| D-Curve | 720 | 599 | 2373 | 1488 |
| U-Curve | 720 | 531 | 1804 | 1097 |
| Overlap | 720 | 521 | 1779 | 422 |

Table S6: Number of metacells in two curves and the overlap between two curves.

Figure S11 shows that the WGCNA algorithm fails to detect gene modules in either the ExDp or ExM-U cell-types, when Pearson's correlation matrices for gene expression is used as input to WGCNA. There are 441 genes and 262 metacells for ExDp and 440 genes for 324 metacells for ExM-U. This result contrasts with the module structure discovered using CSN data as input to PisCES (Figure S10).

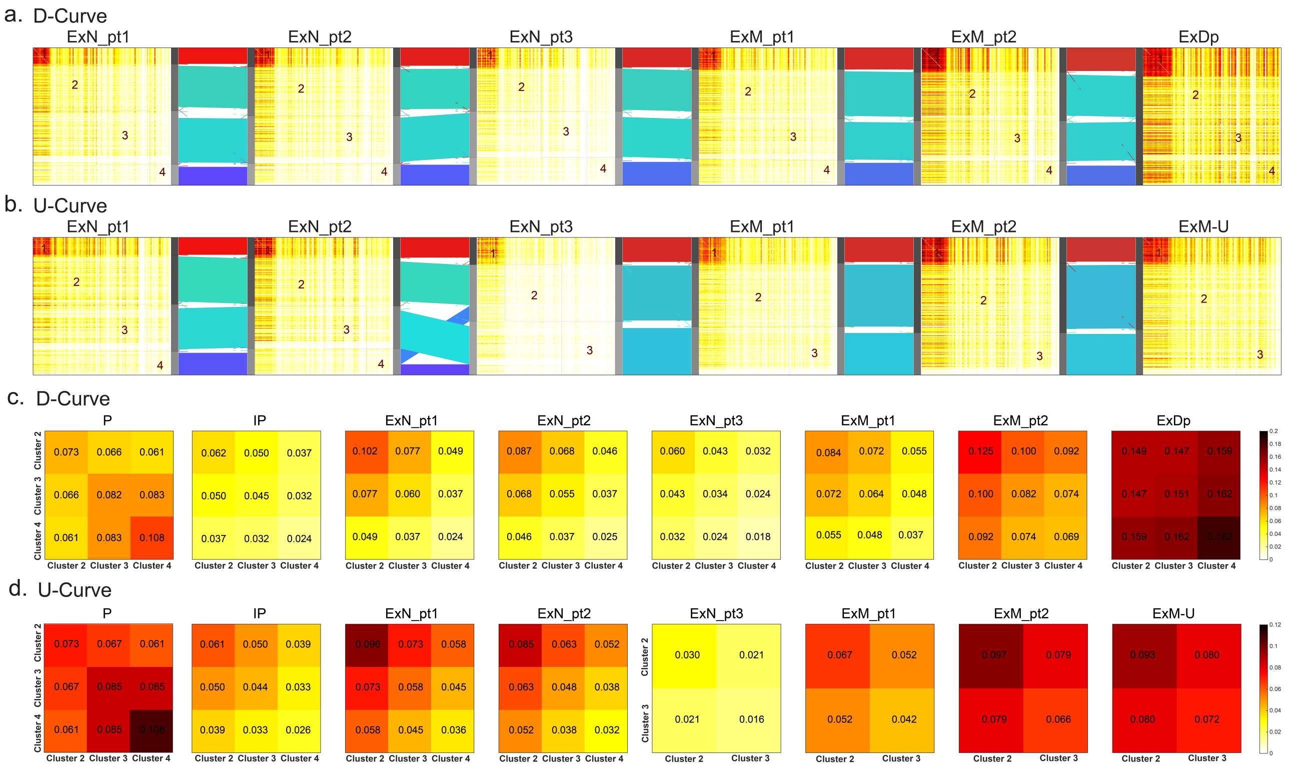


Figure S10: Heatmaps of CSN connections in two curves. Sankey plots of two curves with heatmaps. (**a**) Sankey plots of D-curve with heatmaps, starting from ExN_pt1 to ExDp. (**b**) Sankey plots of U-Curve with heatmaps, starting from ExN_pt1 to ExM-U. (**c** and **d**) Heatmaps of average connection in non-dense clusters. (**c**) D-curve. (**d**) U-curve.

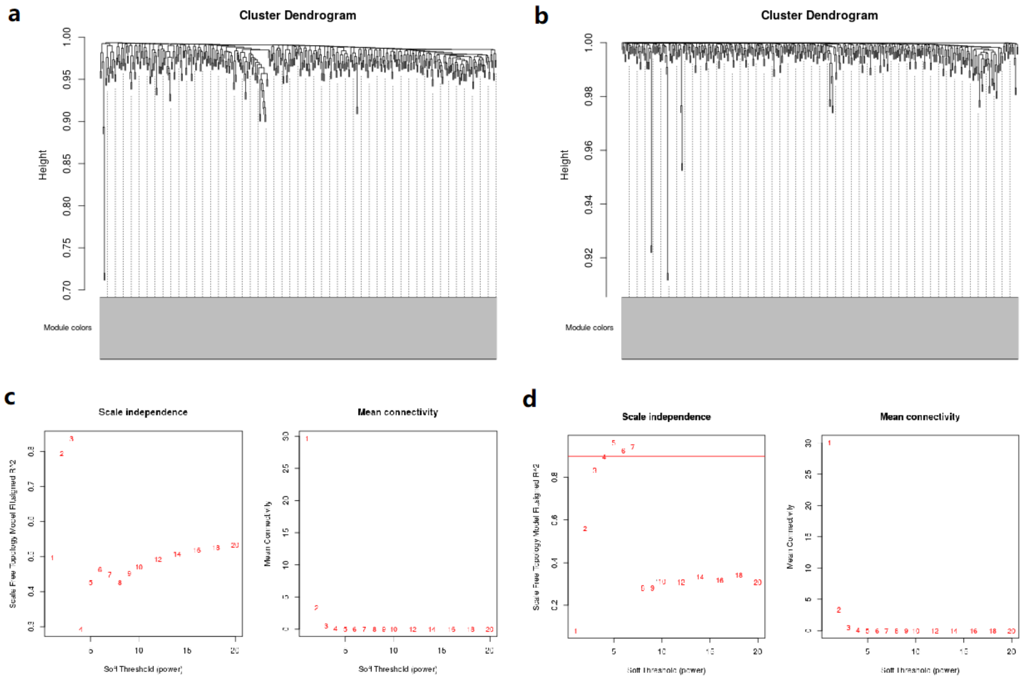


Figure S11: Gene modules generated from Pearson's correlations using WGCNA. (**a**) ExDp from D-curve; (**b**) ExM-U from U-curve. (**c**) power selection plots for ExDp from D-curve: power = 2. (**d**) Power selection plots for ExM-U from U-curve: power = 4.

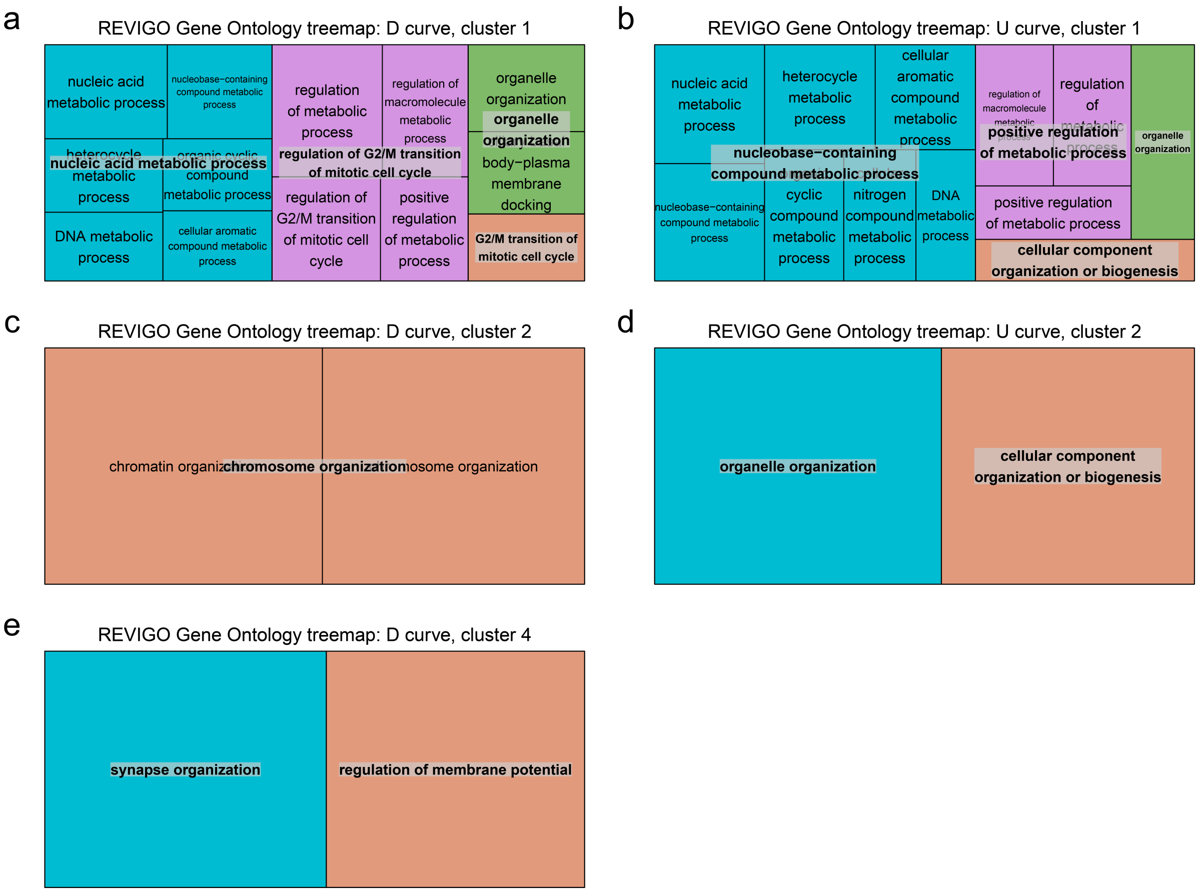


Figure S12: Revigo treemap GO terms for gene clusters in D-curve and U-curve. (**a**) D-curve cluster1 (dense cluster); (**b**) U-curve cluster1 (dense cluster); (**c**) D-curve cluster2; (**d**) U-curve cluster2; (**e**) D-curve cluster4.

#### Supplementary Note 6 CSN analysis of ASD Brain dataset: data processing

The ASD brain dataset [9] consists of single-nuclei RNA-seq measured from 41 samples (22 ASD and 19 controls) from human brains. There are 41,202 expressed genes and 104,559 cells which the authors classify to 17 cell-types: fibrous astrocytes (AST-FB), protoplasmic astrocytes (AST-PP), endothelial (End), parvalbumin interneurons (IN-PV), somatostatin interneurons (IN-SST), SV2C expressing interneurons (IN-SV2C), VIP expressing interneurons (IN-VIP), upper-layer excitatory neurons (L2/3), layer 4 excitatory neurons (L4), deep layer cortico-subcortical excitatory projection neurons (L5/6), Deep-layer cortico-cortical excitatory projection neurons (L5/6-CC), microglia (Mic), immature neurons (Neu-mat), neurogranin expressing neurons I (Neu-NRGN-I), neurogranin expressing neurons II (Neu-NRGN-II), oligodendrocytes (Oligo) and oligodendrocyte precursor cells (OPC) [9]. For our analysis, we merge some cell subtypes together depending on whether the cell-types are distinct in the tSNE plot (Figure 4**a**, Figure S13). For instance, the AST* cell-types are merged into one cell-type, while the L* cell clusters are distinct and analyzed individually (Figure 4**b**).

To circumvent challenges due to sparse counts, which are especially prevalent in single-nuclei RNA-seq data, we cluster similar cells and form metacells [2], which contains around 20 cells each. To avoid batch effects, metacells are created within a sample and cell-type (Table S9). We merged AST-*, IN-*, L* and Neu* together as broad cell-type for the summary of the number of metacells (Table S8). The number of metacells in each cell-type are shown in Table S9. Within a cell-type, some metacells exhibited heterogeneity that was poorly delineated into clusters. For each metacell within a cell-type, we constructed CSNs using the nearby 100 metacells from UMAP plot of the combined ASD and control cells (Figure S14).

For broad cell-type AST, In, L and Neu, more than one original cell-type is included within the broad cell-types. We then determine, based on heterogeneity of cells, whether to analyze the cells within a broad cell-type or within a more refined cell-type. Numbers of metacells in each original cell-type are presented in Table S9. From the UMAP and tSNE plot, we decide to analyze AST as a broad cell-type without division. We divide IN broad cell-type into 2 major cell types (IN-SV2C + IN-VIP) and (IN-SST+IN-PV), the L broad cell type is divided into 4 cell-types: L2/3, L4, L5/6, and L5/6-CC and Neu into 2 cell-types Neu-mat and (Neu-NRGN-I + Neu-NRGN-II) (Figure S13). Cell-types can be analyzed at different levels depending on heterogeneity of the cells and available sample sizes. The original data were partitioned into 17 original cell-types, which spanned 8 broad cell-types. Based on separation of clusters, we performed our analysis on a compromise partition resulting in 13 cell groups, which we refer to as cell-types hereafter. The relationship between the various partitions of cell clusters is shown in Table S7. Metacells for cell-types are shown in Table S9.

| Broad cell-type | AST | | End | IN | | | | Oligo | Mic |
|---|---|---|---|---|---|---|---|---|---|
| Cell-type | AST | | End | IN-PV,SST | | IN-VIP,SV2C | | Oligo | Mic |
| Original cell-type | AST-FB | AST-PP | End | IN-PV | IN-SST | IN-SV2C | IN-VIP | Oligo | Mic |
| Broad cell-type | Neu | | | L | | | | OPC | |
| Cell-type | Neu-mat | Neu-NRGN | | L2/3 | L4 | L5/6 | L5/6-CC | OPC | |
| Original cell-type | Neu-mat | Neu-NRGN-I | Neu-NRGN-II | L2/3 | L4 | L5/6 | L5/6-CC | OPC | |

Table S7: Relation between 17 original cell-types, 8 broad cell-types and the 13 cell-types we feature in our analysis.

| **sampleID** | **1** | **2** | **3** | **4*** | **5*** | **6*** | **7** | **8** | **9*** | **10*** | **11*** | **12*** | **13** | **14** | **15*** | **16** | **17*** | **18*** | **19*** | **20** | **21** | **22*** |
|---|---|---|---|---|---|---|---|---|---|---|---|---|---|---|---|---|---|---|---|---|---|---|
| **AST** | 4 | 18 | 13 | 17 | 18 | 24 | 5 | 13 | 12 | 31 | 8 | 15 | 21 | 10 | 21 | 15 | 11 | 17 | 33 | 29 | 16 | 23 |
| **End** | 1 | 10 | 9 | 3 | 5 | 4 | 4 | 4 | 4 | 3 | 2 | 7 | 23 | 8 | 1 | 2 | 1 | 5 | 5 | 1 | 11 | 5 |
| **IN** | 13 | 36 | 30 | 33 | 26 | 23 | 29 | 11 | 15 | 27 | 14 | 22 | 60 | 19 | 7 | 19 | 17 | 16 | 34 | 27 | 27 | 38 |
| **L** | 9 | 58 | 69 | 49 | 50 | 22 | 15 | 22 | 30 | 55 | 25 | 39 | 148 | 61 | 14 | 36 | 40 | 15 | 45 | 44 | 48 | 65 |
| **Mic** | 2 | 2 | 6 | 2 | 1 | 11 | 2 | 5 | 2 | 9 | 9 | 8 | 9 | 0 | 2 | 10 | 3 | 4 | 7 | 5 | 9 | 2 |
| **Neu** | 13 | 43 | 34 | 69 | 65 | 8 | 26 | 50 | 25 | 14 | 29 | 33 | 100 | 24 | 5 | 29 | 15 | 13 | 28 | 19 | 133 | 33 |
| **Oligo** | 11 | 28 | 54 | 40 | 7 | 24 | 1 | 66 | 19 | 5 | 28 | 22 | 21 | 18 | 5 | 16 | 27 | 3 | 7 | 51 | 17 | 8 |
| **Opc** | 7 | 18 | 20 | 15 | 17 | 11 | 9 | 9 | 6 | 22 | 10 | 10 | 39 | 8 | 13 | 18 | 17 | 8 | 17 | 14 | 11 | 19 |
| **sampleID** | **23*** | **24** | **25*** | **26*** | **27** | **28** | **29** | **30** | **31*** | **32*** | **33*** | **34** | **35** | **36** | **37*** | **38*** | **39** | **40*** | **41*** | | **Ctl** | **ASD** |
| **AST** | 18 | 29 | 32 | 17 | 14 | 14 | 14 | 25 | 17 | 28 | 26 | 6 | 6 | 5 | 16 | 5 | 21 | 17 | 19 | **AST** | 278 | 425 |
| **End** | 3 | 4 | 8 | 1 | 4 | 11 | 3 | 1 | 3 | 4 | 3 | 1 | 2 | 1 | 4 | 2 | 2 | 1 | 1 | **End** | 102 | 75 |
| **In** | 27 | 21 | 40 | 21 | 16 | 31 | 19 | 18 | 18 | 32 | 18 | 26 | 25 | 17 | 33 | 11 | 27 | 6 | 13 | **In** | 471 | 491 |
| **L** | 47 | 58 | 84 | 46 | 30 | 42 | 44 | 22 | 24 | 85 | 49 | 36 | 48 | 19 | 30 | 20 | 33 | 39 | 63 | **L** | 842 | 936 |
| **Mic** | 3 | 9 | 15 | 2 | 13 | 8 | 9 | 9 | 0 | 3 | 7 | 3 | 2 | 0 | 2 | 0 | 2 | 1 | 1 | **Mic** | 105 | 94 |
| **Neu** | 29 | 62 | 75 | 28 | 13 | 40 | 20 | 17 | 21 | 27 | 10 | 28 | 22 | 18 | 19 | 8 | 13 | 16 | 27 | **Neu** | 704 | 597 |
| **Oligo** | 7 | 50 | 49 | 5 | 16 | 8 | 72 | 14 | 1 | 8 | 23 | 45 | 13 | 1 | 3 | 7 | 2 | 9 | 3 | **Oligo** | 504 | 310 |
| **Opc** | 14 | 12 | 26 | 9 | 10 | 8 | 12 | 14 | 11 | 12 | 13 | 16 | 9 | 8 | 16 | 9 | 19 | 14 | 11 | **Opc** | 261 | 300 |

Table S8: The number of metacells for samples and broad cell-types. * indicates the ASD sample while others are the control samples.

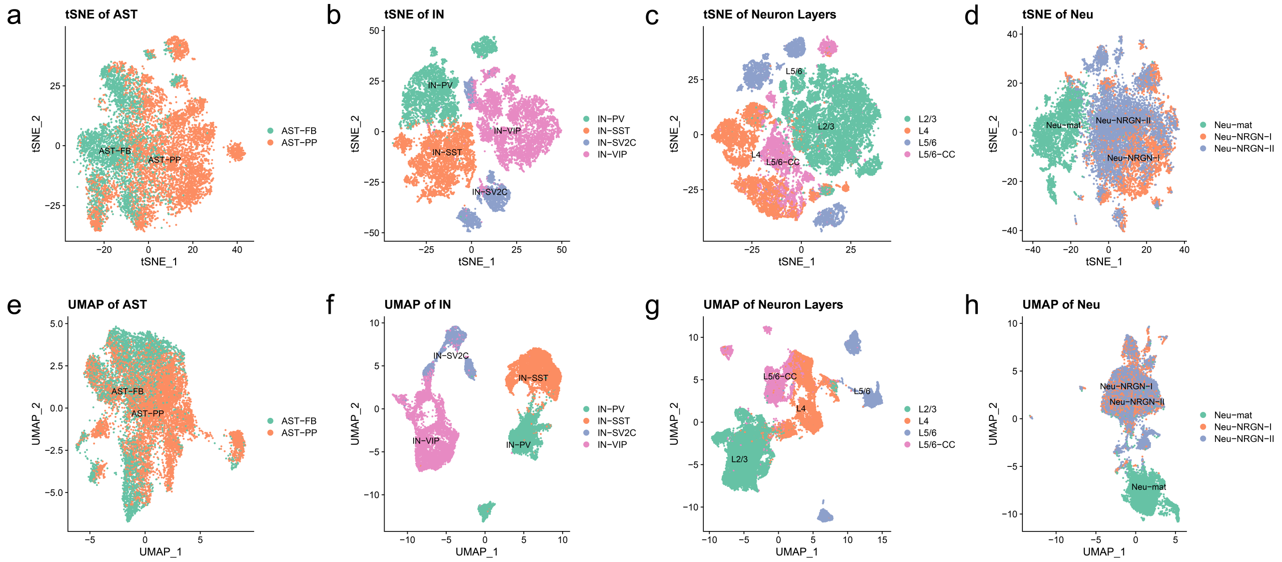


Figure S13: Dimension reduction of 4 broad cell-types, colored by the author defined cell-type labels. (**a**) astrocytes(AST); (**b**) interneurons(IN); (**c**) neuron layers(L); and (**d**) neurons(Neu);

| #metacell | AST | End | IN-VIP,SV2C | IN-PV,SST | L2/3 | L4 | L5/6 | L5/6-CC | Mic | Neu-mat | Neu-NRGN | Oligo | OPC |
|---|---|---|---|---|---|---|---|---|---|---|---|---|---|
| Control | 278 | 102 | 238 | 253 | 414 | 238 | 107 | 177 | 105 | 244 | 353 | 504 | 261 |
| ASD | 425 | 75 | 265 | 206 | 358 | 211 | 109 | 164 | 94 | 235 | 469 | 310 | 300 |
| Total | 703 | 177 | 503 | 459 | 772 | 449 | 216 | 341 | 199 | 479 | 822 | 814 | 561 |

Table S9: Number of metacells for 13 cell-types.

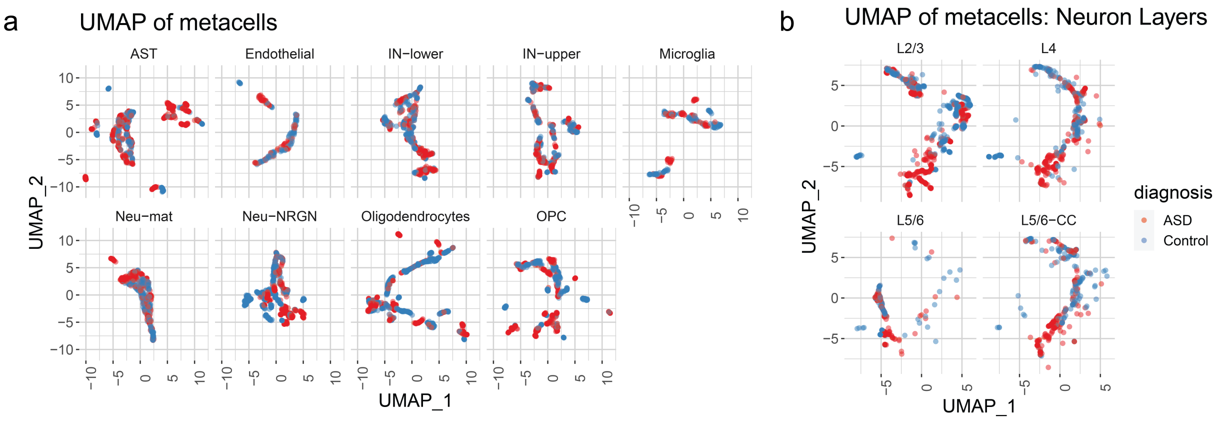


Figure S14: UMAP of metacell expressions for 13 cell-types.

| P-values | AST | End | IN-VIP,SV2C | IN-PV,SST | Mic | Oligogo | OPC |
|---|---|---|---|---|---|---|---|
| sLED-CSN | 0.001* | 0.474 | 0.001* | 0.001* | 0.125 | 0.001* | 0.001* |
| sLED-Pearson | 0.024 | 0.243 | 0.376 | 0.654 | 0.680 | 0.315 | 0.030 |
| DISTp | 0.023 | 0.815 | 0.002 | 0.010 | 0.294 | 0.051 | 0.001* |
| leverage genes | 122 | 103 | 95 | 69 | 83 | 80 | 76 |
| DN genes | 26 | NA | 26 | 26 | NA | 28 | 27 |
| P-values | L2/3 | L4 | L5/6 | L5/6-CC | Neu-mat | Neu-NRGN | |
| sLED-CSN | 0.001* | 0.001* | 0.001* | 0.001* | 0.022 | 0.002* | |
| sLED-Pearson | 0.089 | 0.861 | 0.564 | 0.348 | 0.341 | 0.174 | |
| DISTp | 0.001 | 0.039 | 0.001* | 0.001* | 0.256 | 0.002* | |
| leverage genes | 79 | 112 | 89 | 87 | 146 | 110 | |
| DN genes | 27 | 26 | 24 | 27 | NA | 26 | |

Table S10: p-values from sLED-CSN, sLED-Pearson and DISTp for all 13 cell-types. * indicates significant difference (p-value $< 0.0038$) after adjusted for multiple testing. The leverage genes are the non-zero entries of the sparse leading eigenvector. We only provide DN genes for significant cell-types, corresponding to genes that explain 90% of the variability among the leverage genes.

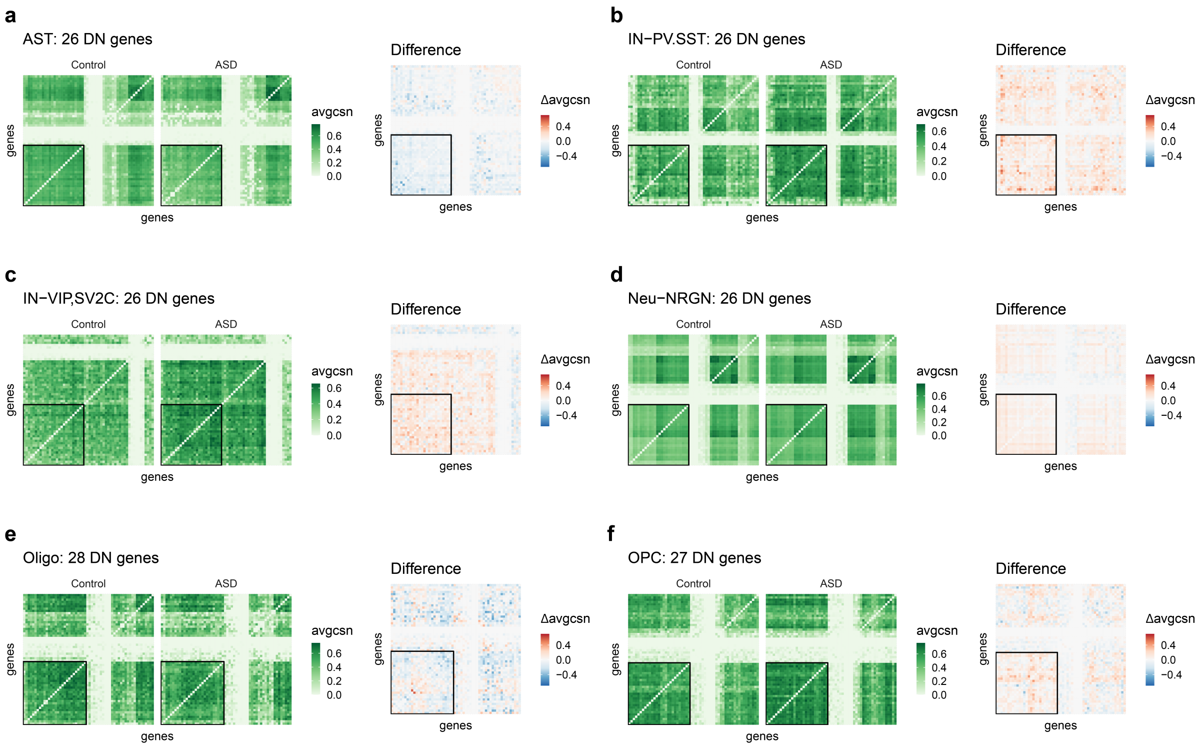


Figure S15: Heatmaps of average CSNs and difference of average CSN between Control and ASD samples. The heatmaps display sLED-CSN DN genes and an additional 30 randomly selected genes from 942 ASD genes. Genes are ordered for each cell-type for display. The DN genes are outlined in black. The green heatmaps show the averaged CSN for control and ASD groups and the red/blue heatmaps show the difference between averaged CSN between control and ASD groups (ASD group minus Control group). (**a**) AST; (**b**) IN-PV,SST; (**c**) IN-VIP,SV2C; (**d**) Neu-NRGN; (**e**) Oligodendrocytes; (**f**) OPC.

In general, DN genes tend to have relatively large variance and DE genes have relatively large mean; and for some cell-types ASD samples are significantly more variable than control samples.

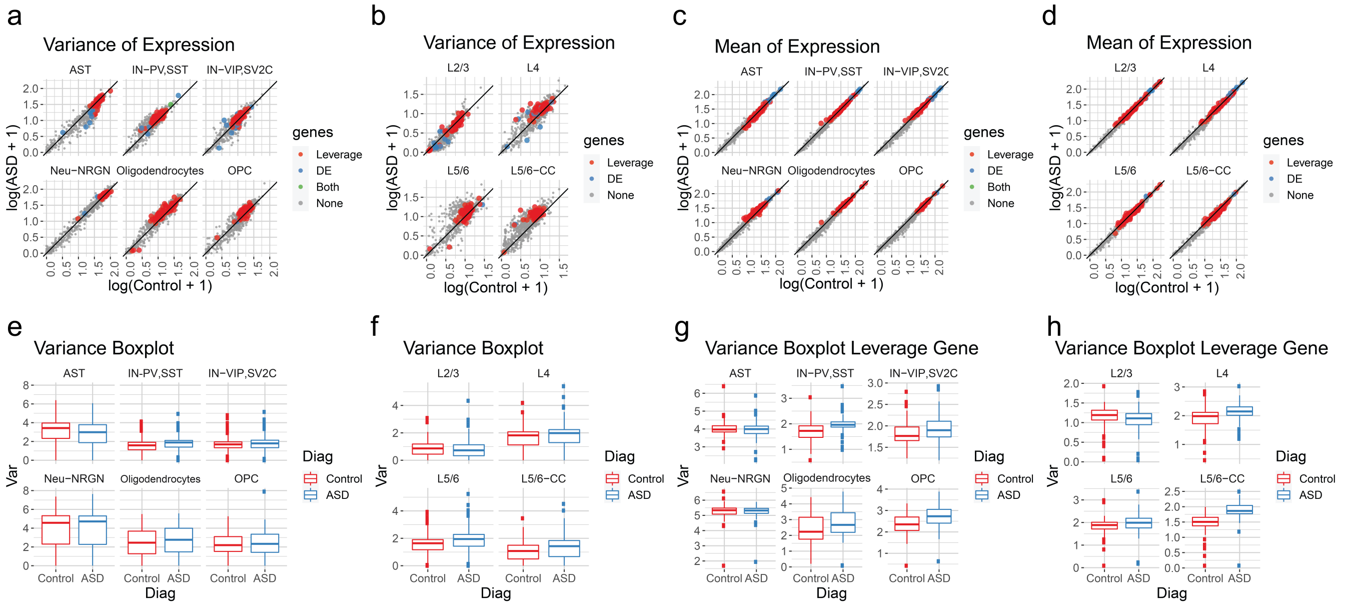


Figure S16: Variance and mean of gene expression of the 942 SFARI genes in the 10 cell-types with significant sLED-CSN signal. (**a-b**) Scatter plots of the variance and mean of gene expression, with x-axis showing the log-transformed values from control group and y-axis from ASD group. The genes are displayed using red and blue to indicate differential network (DN) genes and differential expressed (DE) genes, respectively. (**c**) Boxplots of the variance of gene expression for 942 genes, with red and blue denoting control and ASD, respectively. (**d**) shows the same information for DN genes only.

| Cell-type | AST | IN-PV,SST | IN-VIP,SV2C | Neu-NRGN | Oligo | OPC | L2/3 | L4 | L5/6 | L5/6-CC |
|---|---|---|---|---|---|---|---|---|---|---|
| DN | 26 | 26 | 26 | 26 | 28 | 27 | 27 | 26 | 24 | 27 |
| P-value | 0.074 | 0.023 | 0.006 | 0.474 | 0.109 | 0.001* | 0.001* | 0.050 | 0.012 | 0.040 |

Table S11: p-values from sLED-CSN after removing DN genes. The removal is for the 10 cell-types with significant signal in the original analysis (Table S10). * indicates significant difference (p-value < 0.005) after adjustment for multiple testing.

#### Supplementary Note 7 Gene Ontology (GO) term treemap

The Gene Ontology (GO)[1] describes our knowledge of the biological domain with respect to three aspects: Molecular function. Cellular component and Biological process. In this paper, we focus on biological process. The p-values for GO terms indicate enrichment of the selected gene list in a GO category. Using all ASD genes as the gene universe, an FDR adjusted $p < 0.01$ was considered to be statistically significant. GO treemaps are created by REVIGO with default setting [7] and the areas in GO treemaps indicate the absolute log10 p-value of GO terms.

#### Supplementary Note 8 Runtime of locCSN

The runtime of locCSN are provided below. Three different settings are simulated by ESCO [8] with different sizes of expression matrices (Table S12). For a large set of genes, for example Setting 3 with 1000 genes, parallel computing is recommended to speed up the process of generating CSNs. We also speed up our algorithm by an approximate CSN calculation, which partitions the outcome space for each pair of genes into a grid. Cells that fall into the same grid yield the same test statistic (called fuzzy). With these approximations CSN can be readily applied to very large datasets with good accuracy Figure S17.

| | Setting 1 | Setting 2 | Setting 3 |
|---|---|---|---|
| Number of genes | 100 | 100 | 1000 |
| Number of cells | 200 | 500 | 200 |
| Number of expressed entries | 13114 | 31065 | 39572 |
| Runtime | 322.59s | 1020.744s | NA |
| Speed-up | 186.96s (fuzzy) | 691.62s (fuzzy) | 2260.15s (parallel) |

Table S12: Runtime of locCSN. Python 3.7.6 [MSC v.1916 32 bit (Intel)]

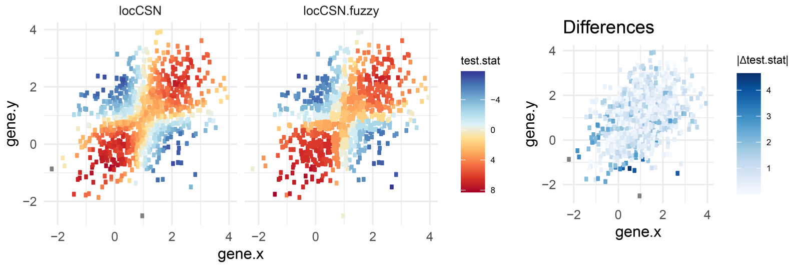


Figure S17: Comparison of test statistics from locCSN and its grid based (fuzzy) approximation. With the same simulation setting in Figure S1**a**, we simulate bivariate normal distribution with $\rho = 0.4$. The left panel is colored by test statistics calculated from locCSN while the right panel is colored by test statistics from locCSN fuzzy approximate.
